# Anatomical and physiological foundations of cerebello-hippocampal interactions

**DOI:** 10.1101/403394

**Authors:** TC Watson, P Obiang, A Torres-Herraez, A Wattilliaux, P Coulon, C Rochefort, L Rondi-Reig

## Abstract

Multiple lines of evidence suggest that functionally intact cerebello-hippocampal interactions are required for appropriate spatial processing. However, how the cerebellum anatomically and physiologically engages with the hippocampus to sustain such interactions remains unknown. Using rabies virus as retrograde transneuronal tracer, we reveal that the dorsal hippocampus receives input from topographically restricted and disparate regions of the cerebellum. By simultaneously recording local field potential from both the dorsal hippocampus and anatomically connected cerebellar regions, we additionally demonstrate that the two structures interact, in a behaviorally dynamic manner, through subregion-specific synchronization of neuronal oscillations in the 6-12Hz frequency range. Together, these results reveal a novel neural network macro-architecture through which we can understand how a brain region classically associated with motor control, the cerebellum, may influence hippocampal neuronal activity and related functions, such as spatial navigation.

## Introduction

The cerebellum is classically associated with motor control. However, accumulating evidence suggests its functions may extend to cognitive processes including navigation [1–6]. Indeed, anatomical and functional connectivity has been described between cerebellum and cortical areas that are engaged in cognitive tasks [7–12]. Furthermore, the cerebellum has recently been found to form functional networks with subcortical structures associated with higher-order functions, such as the basal ganglia [13], ventral tegmental area [14] and hippocampus [15–18].

In the hippocampus, spontaneous local field potential (LFP) activity [19–22] and place cell properties [23], are profoundly modulated following cerebellar manipulation [24 for review]. A recent study has also described, at the single cell and blood-oxygen-level-dependent signal level, sustained activation in the dorsal hippocampus during optogenetic enhancement of cerebellar nuclei output in head-fixed mice [25]. These data point towards the existence of an anatomical projection from the cerebellum to the hippocampus. The suggestion of a direct connection between these two structures has been further supported by a recent tractography study in humans [26] and the presence of short-latency evoked field potentials (2-4 ms) in cat and rat hippocampi after electrical stimulation of the cerebellar vermal and paravermal regions [27–31]. However, secondary hippocampal field responses have also been described, at a latency of 12-15ms following cerebellar stimulation, suggesting the existence of an indirect pathway [28].

Taken together, these studies provide compelling physiological evidence of cerebellar influences on the hippocampus. Yet, they do not provide direct evidence of neuroanatomical connectivity between the two regions. Given the known complex, modular functional and anatomical organization of the cerebellum [32] this represents a major gap in our understanding of the network architecture linking the two structures. What’s more, these studies provide no direct measure of physiological interactions between the regions, which are thought to be essential for maintaining distributed network functions [e.g. 33].

Therefore, this study addresses two fundamental, unanswered questions: which regions of the cerebellum are anatomically connected to the hippocampus and what are the spatio-temporal dynamics of cerebello-hippocampal interactions during behavior? To address these unresolved questions, we used rabies virus as a retrograde transneuronal tracer to determine the extent and topographic organization of cerebellar input to the hippocampus. Based upon the anatomical tracing results, we then studied interactions between the two structures by simultaneously recording LFP from both the cerebellum and the dorsal hippocampus in freely-moving mice. We reveal that specific cerebellar modules are anatomically connected to the hippocampus and that these inter-connected regions dynamically interact during behavior.

## Results

To study the topographical organization of ascending, cerebello-hippocampal projections, we unilaterally injected rabies virus (RABV), together with cholera toxin β-subunit (CTb), into the left hippocampal dentate gyrus (DG). The use of CTb allowed us to identify and measure the extent of the injection sites (Figure S1).

### A precise topography of the cerebellum regions projecting to the hippocampus

We characterized the presence of retrograde transneuronally RABV-infected neurons after survival times of 30, 48, 58 and 66 h [34,35]. Importantly, we did not find any RABV or CTb labeling in the cerebellum at 30h post infection (p.i.), ruling out the existence of a direct cerebello-hippocampal DG pathway in mice. Rather, RABV/CTb-labeled neurons were found in two well described subcortical pathways leading to the DG of the hippocampus (first cycle of infection). One labeled pathway included the diagonal band of Broca and the septum. The other labeled hippocampal input pathway included the lateral entorhinal and perirhinal cortices (Figure S2)[36–38]. At 48 h p.i., a few weakly RABV+/CTb-neurons were found in contralateral deep cerebellar and vestibular nuclei (Fig. 1A, inset), likely reflecting the onset of a second infection cycle. In agreement with this hypothesis, after 58 h p.i., i.e. inside the 12 h time window required for completion of a viral replication cycle [39] we found robust RABV labeling bilaterally in fastigial, dentate and vestibular nuclei, and a small number of weakly labeled neurons in the posterior nucleus interpositus (Fig. 1B-F). At 58 h p.i. we also observed few labeled cells in the cerebellar cortex (Fig. 1A, inset), suggesting the beginning of an overlapping, third infection cycle. Within the two most labeled cerebellar nuclei, fastigial and dentate, RABV-labeled cells were found to be topographically restricted to caudal and central regions, respectively (Fig. 1G).

**Figure 1.**
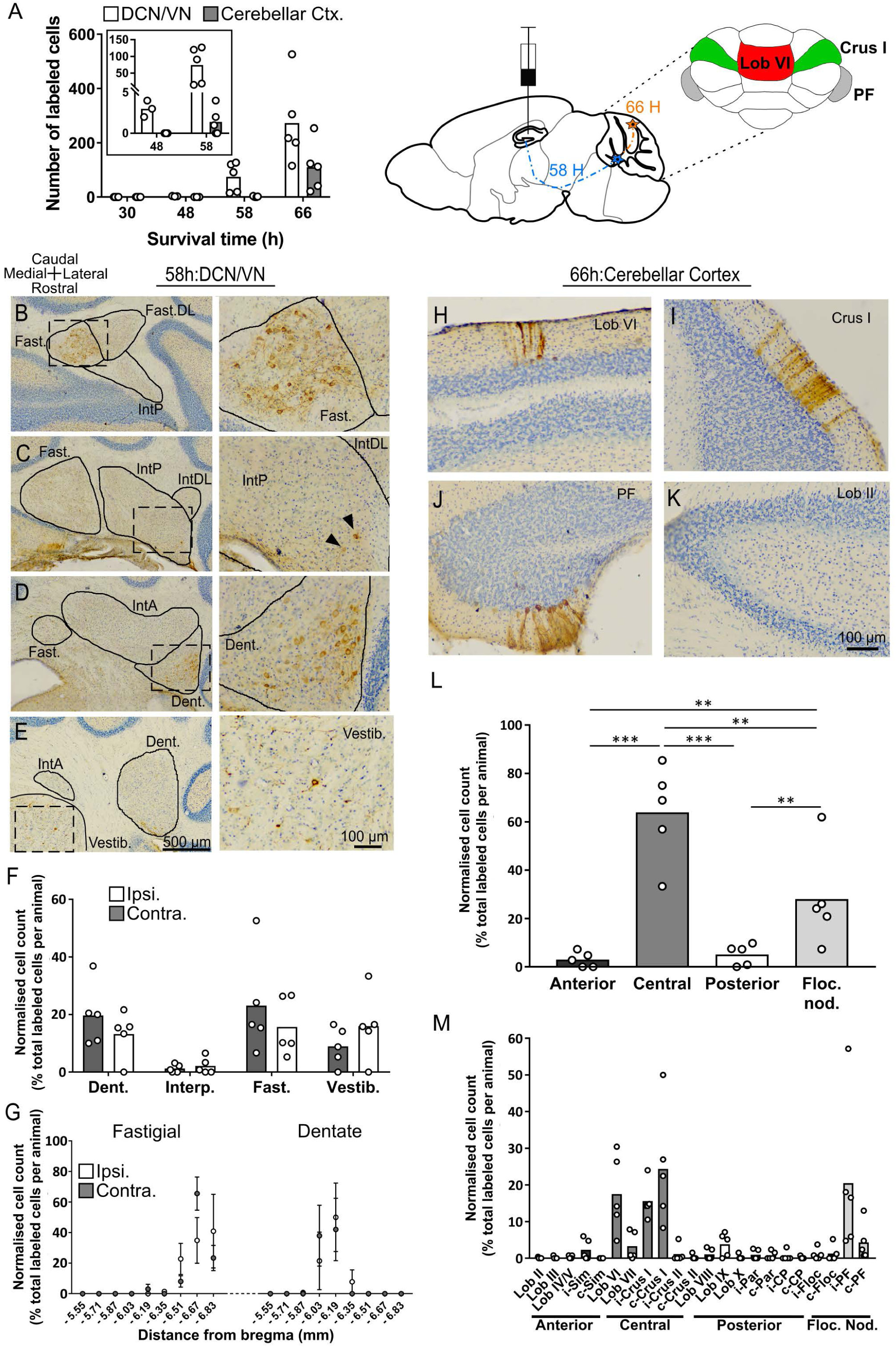
Topographically restricted regions of cerebellar cortex are connected to the hippocampus. **A**, Left, mean number of labeled cells in the deep cerebellar nuclei (DCN), vestibular nuclei (VN) and cerebellar cortex at different survival times following rabies injection in left hippocampal dentate gyrus. Box shows a magnification of the labeling at 48 and 58 h (n = 5 mice for all data points except 48 h, n = 3). Middle, schematic representation of rabies injection site and survival times required to reach the cerebellar and vestibular nuclei (58 h, dashed blue line, polysynaptic pathway), and cerebellar cortex (66 h, orange line). Upper right, schematic view of the posterior cerebellar cortex indicating regions of highest labeling following rabies virus injection (red, vermis lobule VI; green, Crus I; grey, paraflocculus). **B-E**, Representative photomicrographs showing labeling in the contralateral cerebellar and vestibular nuclei 58 h post infection. Left panels show low magnification view, right panels show magnified view of area indicated by dashed box. Solid arrow heads indicate the presence of the very few labeled cells in the IntP. **F**, Pooled, normalized counts of rabies labeled cells in the ipsi-and contralateral cerebellar and vestibular nuclei 58 h post infection (n = 5 mice). No significant differences were found between ipsi-and contralateral nuclei (nuclei x hemisphere two-way ANOVA, hemisphere effect F _(1,4)_ = 1.31×10^−5^, p = 0.99, interaction effect F _(3,12)_ = 2.79, p = 0.09, nuclei effect F _(3,12)_ = 9.38, p = 0.002). **G**, Normalised cell counts in the fastigial nucleus (left) and dentate nucleus (right), according to their rostro-caudal position relative to bregma. Open circles, contralateral count; filled circles, ipsilateral count (n= 5 mice). **H-K**, Representative photomicrographs of the resultant labeling in lobule VI, Crus I, paraflocculus and lobule II at 66 h post infection. **L**, Normalised count of rabies labeled cells in anterior (black bar; lobule II to lobule IV/V); central (dark grey bar; lobule VI to Crus II); posterior (clear bar; lobule VIII and lobule IX) and flocculonodular (Floc. Nod., light grey bar; lobule X, flocculus and paraflocculus) cerebellum 66 h post infection (n= 5 mice; one-way ANOVA with FDR correction, F _(3,16)_ = 19.11, p < 0.0001). **M**, Normalised cell count of rabies labelled cells in all assessed lobules 66 h post infection. Colour coding of bars indicate assignment of lobules to either anterior, central, posterior or vestibular cerebellum as indicated in **L**. Abbreviations: Dent., Dentate nucleus; Fast., fastigial nucleus; Fast. DL, dorsolateral fastigial nucleus; Floc. Nod., flocculonodular lobe; Interp., nucleus interpositus; IntA, nucleus interpositus anterior; IntDL, dorsolateral nucleus interpositus; IntP, nucleus interpositus posterior; i-Sim, ipsilateral simplex lobule; c-Sim, contralateral simplex lobule; i-Crus I, ipsilateral Crus I; c-Crus I, contralateral Crus I; i-Crus II, ipsilateral Crus II; c-Crus II, contralateral Crus II; i-Par, ipsilateral paramedian lobule; c-Par, contralateral paramedian lobule; i-CP, ipsilateral copula; c-CP, contralateral copula; i-Floc, ipsilateral flocculus; c-Floc, contralateral flocculus; i-PF, ipsilateral paraflocculus; c-PF, contralateral paraflocculus; Vestib., vestibular nuclei. ** q < 0.01, *** q < 0.001.

Following 66 h of incubation, the number of strongly labeled cells increased in the DCN and vestibular nuclei (Fig. 1 A); however, the topographical distribution remained unchanged (Fig. S3). At the level of the cerebellar cortex, longitudinal clusters of RABV+ Purkinje cells (PCs) were found in a bilateral manner across highly restricted central and flocculo-nodular regions (Fig. 1L). The bilateral cerebellar patterning of RV labeled cells observed after unilateral hippocampal injections likely reflects the existence of commissural connections within the pathway rather than the existence of bilateral projections arising from the cerebellum. The presence of labeling in the contralateral hippocampus at the earliest survival time (30h; Table 1) and the appearance at 48h of RV labeled neurons exclusively in the contralateral cerebellar output nuclei are consistent with this hypothesis. In the central cerebellum, clusters were particularly concentrated in lobule VI and Crus I (Fig. 1 H, I, M). In the flocculo-nodular cerebellum, RABV-labeled cells were found in the dorsal and ventral paraflocculus (Fig. 1 J and M and Fig. 2B). Within the vermis we identified a single cluster of RABV+ Purkinje cells that extended across both lobule VIa and lobule VIb-c (Fig. 2). In contrast, within Crus I, RABV-labeled Purkinje cells were arranged in two spatially isolated clusters, one located rostro-laterally and the other caudo-medially (Fig. 2).

**Table 1.**
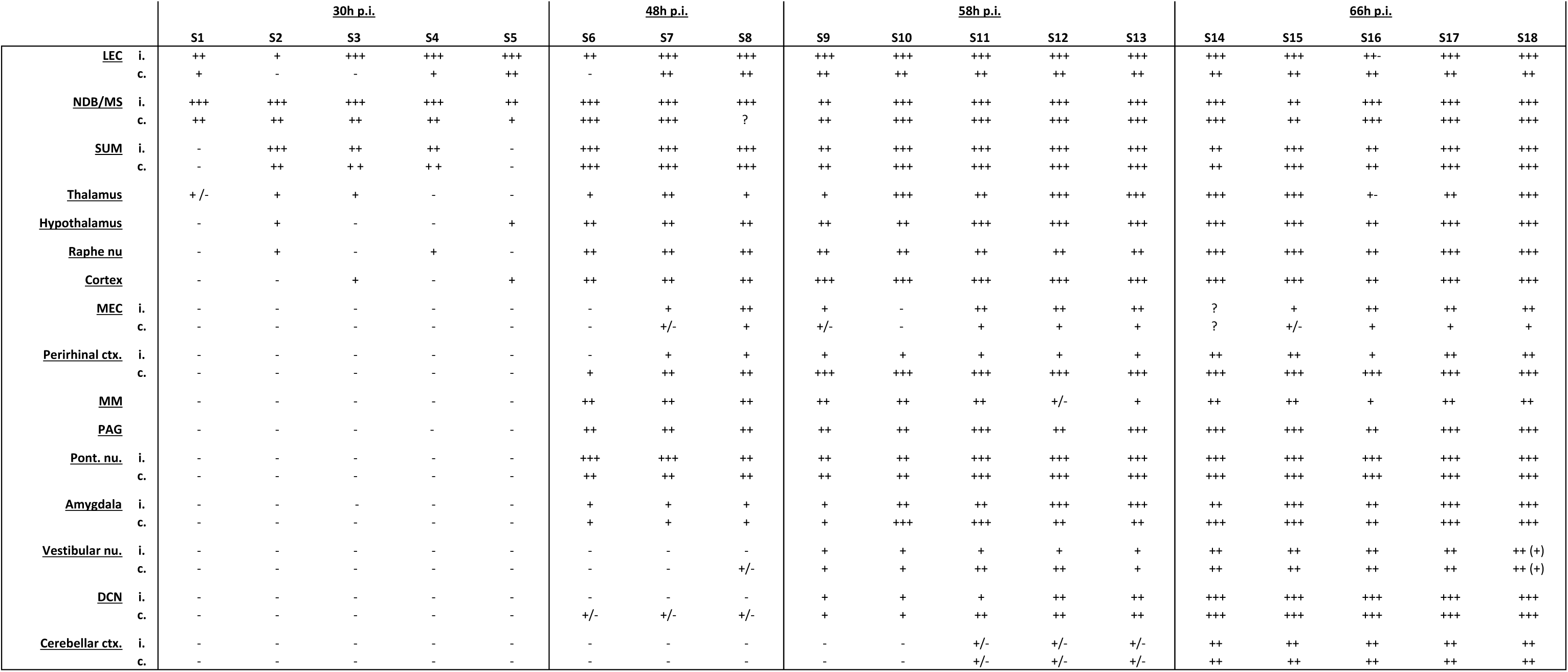
Overview of RABV labeling intensity in different brain regions for all animals in the four experimental groups (30h, 48h, 58h and 66h post RABV injection). (-) denotes no labeling, (+/-) few cells, (+) minor labeling, (++) fair labeling, (+++) strong labelling. Question mark indicates that the area was not available for analysis. When the labeling was different between ipsilateral (i.) and contralateral (c.) hemispheres, the regions are split in two columns.

**Figure 2.**
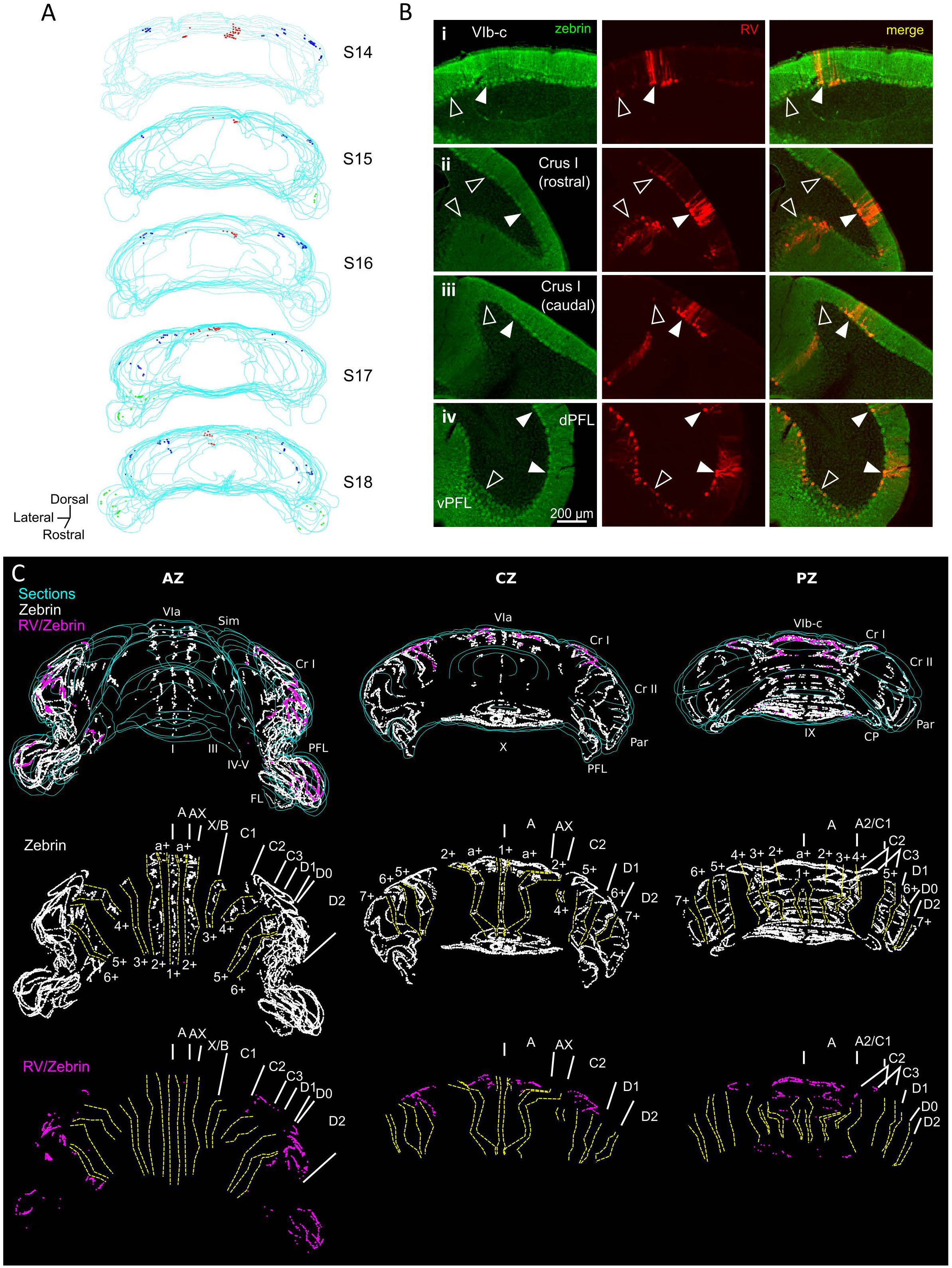
Different cerebellar modules project to the hippocampus. **A**, 3-D reconstruction showing the location of RABV+ Purkinje cells in the most labeled cerebellar lobules at 66 h post-infection. Red, blue and green dots represent RABV+ Purkinje cells in lobule VI, Crus I and paraflocculus, respectively. **B**, Photomicrographs from case S18 showing double staining against zebrin II (green, left column), RABV (red, central column) and merge (right column) in lobule VI (i), Crus I (ii and iii) and paraflocculus (iv). RABV+ Purkinje cells were also zebrin positive and were organized in clusters of strongly labeled RABV+ cells (filled arrow-heads) surrounded by weakly labeled RABV+ Purkinje cells (unfilled arrow-heads). **C**, Assignment of the RABV+ clusters to specific cerebellar modules for case S18 in the anterior (AZ; left), central (CZ; central column) and posterior (PZ; right column) zones. First row shows stacked sections with zebrin positive Purkinje cells (white dots) and RABV+ Purkinje cells, which were also zebrin positive (purple dots, strong and weakly labeled cells included); central row shows reconstructed principal zebrin bands (delineated by yellow dashed lines and named from 1+ to 7+; nomenclature from [42] and cerebellar modules (capital letters; defined as in [42]); and bottom row shows the location of the RABV+/zebrin Purkinje cells (purple dots) in relation to reconstructed zebrin bands and modules. Abbreviations, I, lobule I; III, lobule III; IV/V, lobule IV/V; VIa and VI b-c, lobule VIa and VI b-c, respectively; IX, lobule IX; X, lobule X; Sim, lobule simplex; Cr I, Crus I; Cr II, Crus II; Par, paramedian lobule; CP, copula, PFL, paraflocculus, FL, flocculus.; dPFC and vPFC, dorsal and ventral paraflocculus, respectively.

The topographical arrangement of RABV-labeled PCs in longitudinal clusters is in keeping with the well-described modular organization of the cerebellum [e.g. 32]. Mapping of molecular marker expression patterns, such as zebrin II banding, provides a reliable basis from which modules can be defined and recognized in the cerebellar cortex of rodents. Thus, to further assign the observed PC clusters to previously described cerebellar zones, we used a double immunohistochemical approach to stain for both RABV and aldolase C (zebrin II) in one animal (case S18) 66 h after infection (Fig. 2B) [40,41]. Lobule VI, Crus I and paraflocculus are mostly zebrin positive regions [42] and we found that RABV-labeled Purkinje cells co-localized with zebrin II in all the observed clusters (Fig. 2B). In the vermis, lobule VIa RABV-labeled PCs were mostly located in the a+ band. The few RABV-labeled cells found in lobule VII were confined to the 2+ band. Thus, together, these labeled cells belong to the a+//2+ pair that constitutes part of the cerebellar A module (Fig. 2C) [43]. In Crus I, the rostrolateral cluster of RABV-labeled PCs was aligned with the anterior 6+ zebrin band corresponding to module D2. The caudomedial cluster was in continuation with the posterior 5+ zebrin band suggesting that it is part of the paravermal module C2 (Fig. 2C). In the paraflocculus, the assignment of the RABV-labeled cells to specific modules was not addressed given the complex morphology of this region. However, the presence of RABV-labeled cells both in the dorsal and ventral paraflocculus suggests the involvement of more than one module (Fig. 2B-C) [44].

Cerebellar modules are also defined by their outputs through the deep cerebellar and vestibular nuclei [32,45]. The presence of RABV-labeled cells in the fastigial nucleus is consistent with the involvement of module A. Similarly, the D2 module is routed through the dentate nuclei in which we find robust RABV labeling. We also found RABV+ cells in the nucleus interpositus posterior, which provides the output of module C2. Finally, RABV labeling was observed in the vestibular nuclei, which may represent the output of RABV+ Purkinje cells clusters observed in the ventral paraflocculus. Together, our neuroanatomical tracing data indicate that cerebellar projections to the hippocampus emanate from three distinct cerebellar modules subserving diverse functions.

### Cerebello-hippocampal physiological interactions in a familiar home-cage environment

In order to question the potential functional relevance of cerebello-hippocampal anatomical connectivity, we implanted mice (n = 21) with arrays of bipolar LFP recording electrodes in bilateral dorsal hippocampus (HPC) and unilaterally in two highly RABV-labeled regions of the central cerebellum, lobule VI (midline) and Crus I (left hemisphere). For comparison, we also simultaneously recorded LFP from cerebellar regions with minimal RABV labeling (lobule II or lobule III; Fig. 1M; Fig. 3A and B). Data were excluded from further analysis in cases where postmortem histological inspection revealed that electrode positions were off-target Fig. S4).

**Figure 3.**
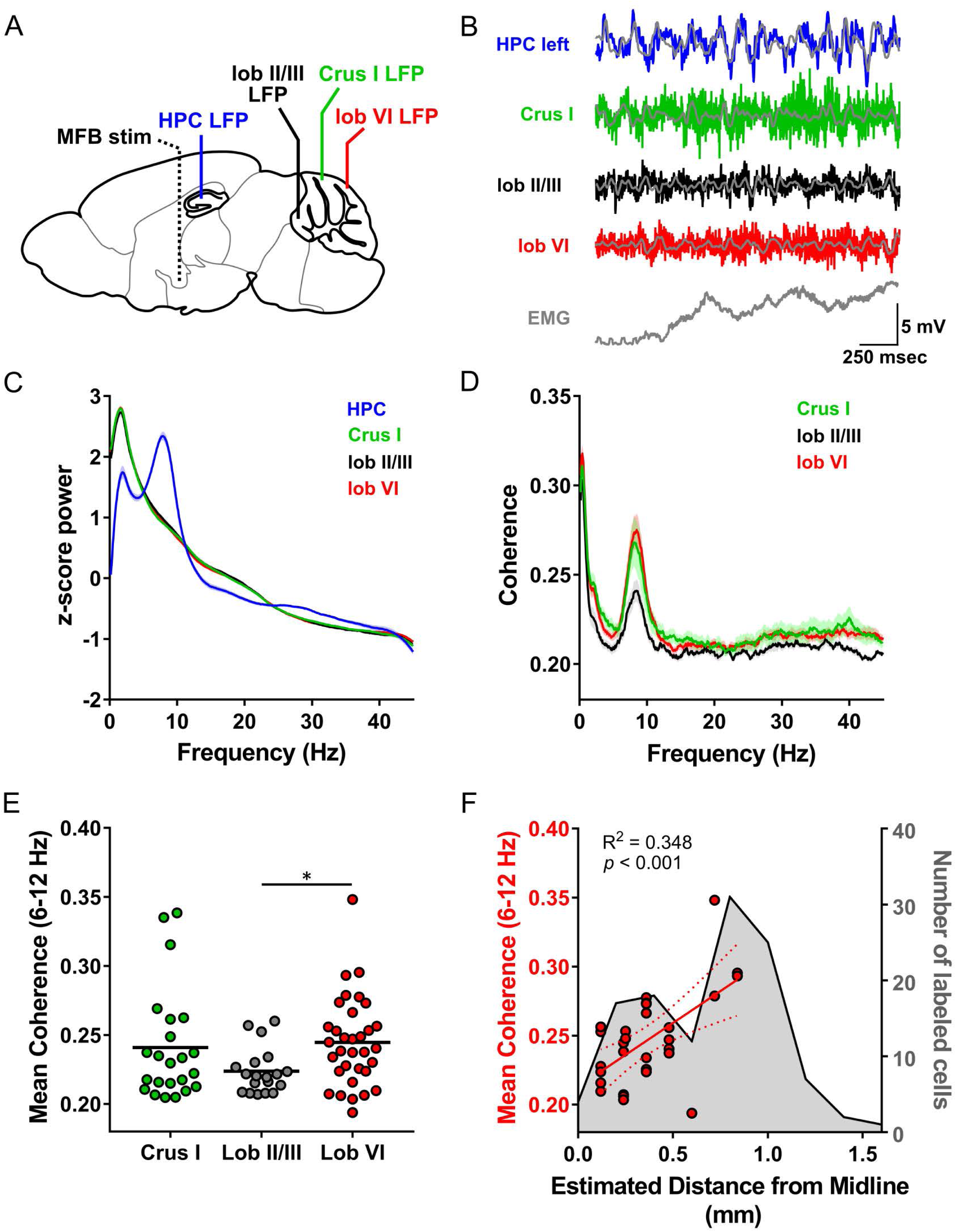
Assessment of cerebello-hippocampal interactions during active movement in the home-cage. **A**, Schematic representation of recording and stimulation electrode implant positions. **B**, Representative simultaneous LFP and EMG recording made during active movement in the homecage condition. Colored lines: raw LFP (filtered from 0.1 to 600 Hz). Overlaid grey lines: LFP filtered from 6-12 Hz. Note prominent 6-12 Hz oscillations in left hippocampal recording and deflections on neck EMG trace, reflecting active movement (EMG rectified and smoothed to 2.5 ms). **C**, Pooled z-score of the power spectra of hippocampal LFP recorded from the left (n = 16 mice) and right (n= 18 mice) hemispheres during homecage exploration and from cerebellar Crus I (n = 12), lobule II/III (n = 10) and lobule VI (n = 18). **D**, Pooled coherence between cerebellar cortical regions (colour coded) and hippocampus (left, Crus I, n = 11; lobule II/III, n = 8; lobule VI, n = 15; right, Crus I, n = 11; lobule II/III, n = 9; lobule VI, n = 16) during homecage exploration. **E**, Mean cerebello-hippocampal coherence in the 6-12 Hz range (Crus I, n = 21 values/12mice; lobule II/III, n = 17 values/10 mice; lobule VI, n = 31 values/18 mice). Lobule VI-hippocampus coherence level was significantly higher than that observed with lobule II/III (*, q < 0.05; Kruskall-Wallis with FDR correction). **F**, Mean 6-12 Hz coherence between lobule VI and hippocampus plotted against estimated medio-lateral recording electrode position in lobule VI (red dots; n = 16 mice, coherence with right and left hippocampus pooled; linear regression, R^2^ = 0.348, *p* < 0.001). In grey, number of RABV+ cells counted across lobule VI, 66 h after injection in the left hippocampus as a function of medio-lateral position (0.2 mm bins; n = 5 mice). Shading indicates S.E.M. Abbreviations, LFP, local field potential; HPC, dorsal hippocampus; lob II/III, lobule II/III; lob VI, lobule VI; EMG, electromyogram.

The spectral profile of cerebellar and hippocampal LFP activity was first assessed during active movement in a familiar home-cage environment (see methods; mean speed, 2.7 ± 0.3 cm/s). Within the HPC, a dominant 6-12Hz theta oscillation was similarly observed in both hemispheres (Fig. S5B; left HPC: peak spectral frequency = 7.81 ± 0.13 Hz, mean 6-12Hz z-score power = 1.54 ± 0.07, N = 17 mice; right HPC: peak spectral frequency = 7.72 ± 0.12 Hz, mean 6-12Hz z-score power = 1.54 ± 0.07, N = 19 mice; unpaired t test, t_34_= 0.007, *p* = 0.99). Fig. 3C shows combined spectra from both left and right HPC peak spectral frequency = 7.76 ± 0.09 Hz, mean 6-12Hz z-score power = 1.55 ± 0.05).

Although a clear peak in the 6-12 Hz band was not detected in cerebellar recordings, transient 6-12 Hz oscillations were recorded (Fig. 3C and S6). The mean 6-12 Hz z-score power did not differ between the different cerebellar recording sites (Fig. 3C; Crus I: 0.80 ± 0.30, N = 13 mice; lobule II/III: 0.84 ± 0.03, N = 11 mice; lobule VI: 0.79 ± 0.02, N = 19 mice; one-way ANOVA, F _(2,40)_ = 0.85, *p* = 0.43).

As an indicator of cross-structure interaction [33], we next calculated coherence between LFP recorded from the different cerebellar subregions and left or right HPC. We found no statistically significant influence of hippocampal laterality on cerebello-hippocampal coherence (Fig. S5C-E hemisphere x combination two-way ANOVA, hemisphere effect F _(1,69)_= 0.23, *p* = 0.64, interaction effect, F_(2,69)_ = 0.06, *p* = 0.94). Therefore, for further analysis, we grouped these coherence values.

A single peak in coherence was observed for all cerebello-hippocampal combinations in the theta frequency range (6-12 Hz, Fig. 3D; Crus I-HPC peak coherence = 7.99 ± 0.13 Hz, lobule II/III-HPC peak coherence = 8.75 ± 0.16 Hz, lobule VI-HPC peak coherence = 8.55 ± 0.11 Hz). However, significant differences across combinations were observed within this bandwidth and LFP oscillations were significantly more synchronised between HPC and lobule VI than with lobule II/III (Fig. 3D and E; mean lobule VI-HPC coherence, 0.245 ± 0.006; mean lobule II/III-HPC coherence, 0.223 ± 0.004; Kruskal-Wallis with FDR correction, *q* = 0.045; lobule VI, n = 33 values/20 mice; lobule II/III, n = 19 values/11 mice). Within lobule VI, coherence was significantly correlated to the mediolateral position of the recording electrode, which was consistent with the mediolateral location of greatest RABV-labeled PCs (Fig. 3F; linear regression, *R*^*2*^ = 0.35, F _(1,27)_ = 14.45, *p* = 0.0007). Mean coherence between HPC and Crus I (0.24 ± 0.01; n = 23 values/13 mice) was not significantly higher than with lobule II/III (Fig. 3E; Kruskal-Wallis with FDR correction, *q* > 0.05).

### Cerebello-hippocampal interactions during the learning of a goal-directed behavior

To further characterize the dynamics of cerebello-hippocampal interactions, we quantified cerebello-hippocampal theta coherence during a goal-directed task. A subset of mice (n=8) were trained to traverse a linear track to get a reward (medial forebrain bundle stimulation, see methods) at a fixed position (Fig. 4A).

**Figure 4.**
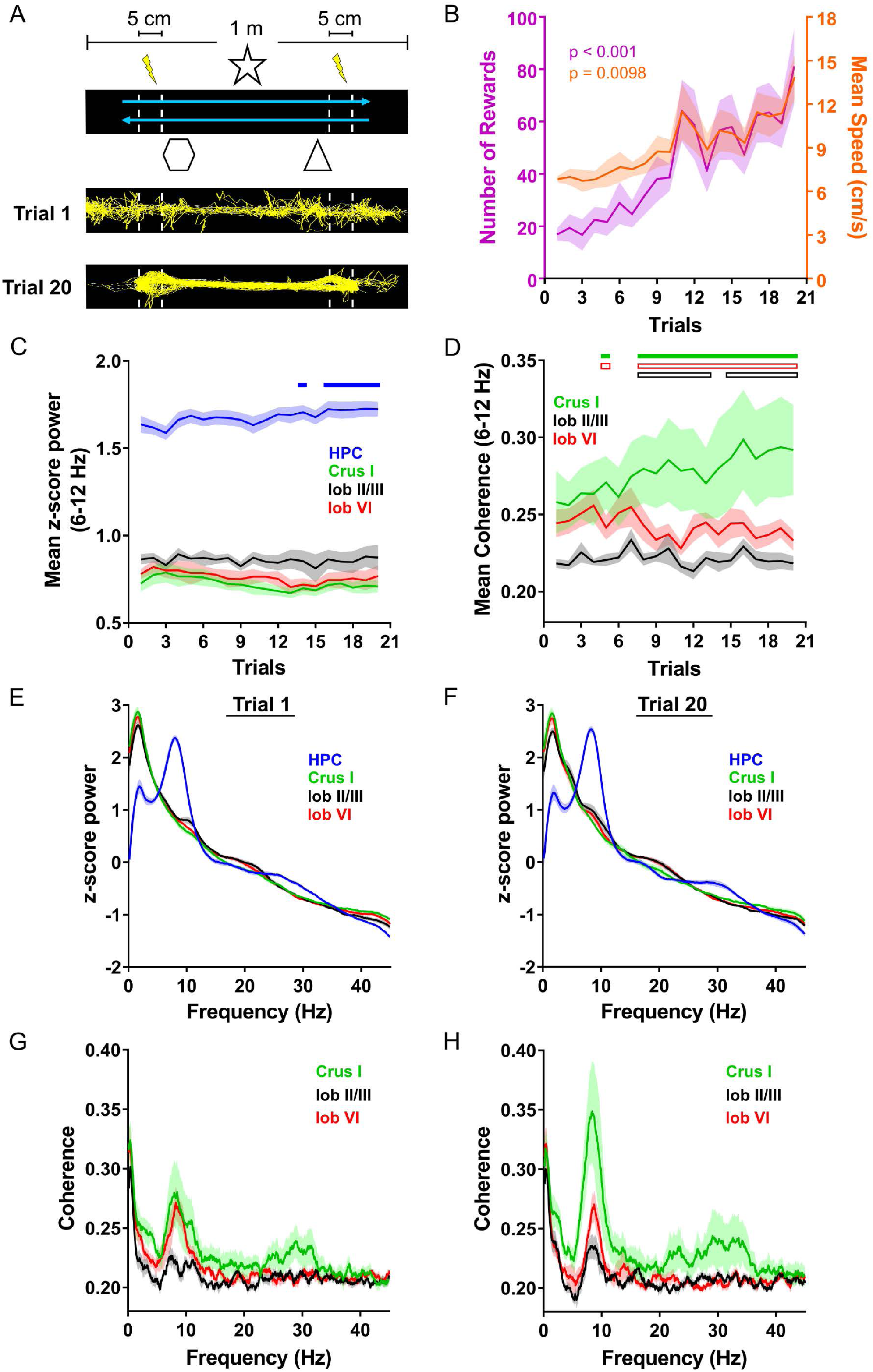
Cerebello-hippocampal interactions during goal-directed behavior A, Mice learned to traverse a 1 m linear track to receive a medial forebrain bundle stimulation upon reaching invisible goal zones (lightening symbols indicate MFB stimulation; n = 8 mice). Representative trajectories from early (trial 1) and late training (trial 20) show the transition from exploratory to goal-directed behavior. **B**, Mice improved their performance in the task across trials as shown by increases in the mean number of rewards obtained (purple line; plotted against left Y axis; one-way repeated measures ANOVA, time effect p < 0.0001) and the mean speed (orange line; plotted against the right Y axis; one-way repeated measures ANOVA, time effect p = 0.0098).**C**, Mean z-score 6-12 Hz power of the recorded LFP signals (colour coded; Crus I, n = 5; lobule II/III, n = 6; lobule VI, n = 7; HPC left, n = 6; HPC right, n = 7; left and right HPC values are pooled as no difference was observed across hemispheres, hemisphere x trial two-way ANOVA with FDR correction, hemisphere effect *p* = 0.5974, interaction effect *p* = 0.3132, trial effect *p* < 0.0001) across trials. Solid blue line indicate trials where hippocampus values were significantly higher than those in trial 1 (q < 0.05).**D**, Mean coherence in the 6-12 Hz range between cerebellar regions (colour coded) and hippocampus (pooled bilaterally) during learning of the linear track task (Crus I, n = 8 values/5 mice; lobule II/III, n = 9 values/6 mice; lobule VI, n= 12 values/7 mice). Significant changes in coherence were observed over trials (combination x trial two-way ANOVA with FDR correction, trial effect p < 0.001, combination effect p = 0.0285, interaction effect p < 0.0001). Solid green rectangles indicates trials where Crus I-hippocampus coherence was higher than those in trial 1 (q < 0.05). This increase in Crus I-hippocampus coherence was also reflected in significant changes relative to other lobules: grey bordered rectangle corresponds to trials significantly higher than lobule II/III-hippocampus coherence (q < 0.05) while red bordered rectangles indicates those significantly higher than lobule VI-hippocampus coherence (q < 0.05). **E-F**, Pooled power spectra from hippocampal and cerebellar LFPs (z-score normalized, colour coded) from representative trials of early (**E**; trial 1) and late (**F**; trial 20) stages of training. **G-H**, Pooled coherence between cerebellar cortical regions and bilateral hippocampus from representative trials of early (**G**; trial 1) and late (**H**; trial 20) stages of training. Shading indicates S.E.M. Abbreviations, HPC, dorsal hippocampus; lob II/III, lobule II/III; lob VI, lobule VI.

Across training, mice improved their performance as shown by the optimisation of their path (Fig. 4A), significant increase in the number of rewards obtained per trial (Fig. 4B; mean number of rewards obtained on 1^st^ trial = 16 ± 3, mean number of rewards obtained on 20^th^ trial = 81 ± 15; one-way repeated measures ANOVA, F_(2.882,20.18)_ = 8.93, *p* < 0.001) and the significant increase in their mean speed (Fig. 4B; mean speed on 1^st^ trial = 6.61 ± 0.30 cm/s, mean speed on 20^th^ trial = 13.86 ± 1.56 cm/s; one-way repeated measures ANOVA, F_(2.45,17.15)_ = 5.631, *p* = 0.0098). Thus, we next explored the dynamics of LFP power and cerebello-hippocampal 6-12Hz coherence across this learning period.

In the hippocampus, theta oscillations remained dominant in the recorded LFP throughout training. A significant increase in both the mean theta power (Fig. 4C, S7A) and the peak frequency (Fig. S7B) was observed in parallel with the performance and this was independent of the hippocampal hemisphere (theta power: trial effect F_(19,209)_ = 3.11, p < 0.0001, hemisphere effect F_(1,11)_ = 0.30, p = 0.60; interaction effect, F_(19,209)_ = 1.14, p = 0.31; peak frequency: trial effect, F_(19,209)_ = 8.84, p < 0.0001, hemisphere effect F_(1,11)_ = 0.73, p = 0.41, interaction effect F_(19,209)_ = 0.16, p > 0.99). In accordance with this finding, post-hoc analysis revealed that mean theta power and peak frequency were significantly different between first and last trials (Fig. 4E-F; mean z-score HPC theta power; mean on 1^st^ trial = 1.64 ± 0.05, mean on 20^th^ trial = 1.72 ± 0.04, q = 0.0180; peak frequency, mean trial 1 = 8.02 ± 0.15 Hz, mean trial 20 = 8.26 ± 0.13 Hz, q = 0.0205).

In the cerebellum, a global variation in the mean theta power was observed across trials but no difference was found between Crus I, lobule VI and lobule II/III and no significant variation was found between the last and first trials (Fig. 4C; Crus I: mean trial 1 theta power = 0.73 ± 0.05, mean trial 20 theta power = 0.71 ± 0.03, N = 5 mice; lobule II/III: mean trial 1 LFP power = 0.86 ± 0.03, mean trial 20 theta power = 0.87 ± 0.07, N = 6 mice; lobule VI: mean trial 1 theta power = 0.78 ± 0.03, mean trial 20 theta power = 0.77 ± 0.06, N = 7 mice; cerebellar region x trial two-way repeated measures ANOVA with FDR correction, cerebellar region effect, F_(2,15)_ = 2.88, p = 0.09, trial effect F_(19,285)_ = 3.08, p < 0.0001, interaction effect F_(38,285)_ = 0.72, p = 0.89, no trial was different from trial 1).

However, as learning progressed, cerebello-hippocampal theta coherence evolved in a non-uniform manner (Fig. 4D; trial x combination two-way repeated measures ANOVA, trial effect F _(19,494)_ = 2.42, p < 0.001, combination effect F _(2,26)_ = 4.09, p = 0.028, interaction effect F _(38,494)_ = 3.43, p < 0.0001). Post-hoc analysis revealed that only Crus I-HPC coherence increased significantly across trials compared to initial values (multiple comparisons against trial 1 with FDR correction; *q* < 0.05 for trials 5 and 7-20; Fig. 4D) and this was independent of hippocampal hemisphere (Fig. S7C). Furthermore, this increase resulted in changes in the differences in coherence observed between cerebello-hippocampal recording combinations. Indeed, while during initial trials no significant inter-regional differences were observed, in later trials, Crus I became significantly more coherent with HPC than lobule II/III and lobule VI (Fig. 4D, multiple comparisons between combinations with FDR correction; Crus I vs lobule II/III *q* < 0.05 for trials 5 and 8-20; Crus I vs lobule VI *q* < 0.05 for trials 8-12 and 14-20). Unlike the observed shift in peak frequency of HPC theta power across trials, the peak frequency of theta coherence remained constant for all the cerebello-hippocampal combinations (Fig. S7D; Crus I-HPC: mean peak frequency = 8.41 ± 0.09 Hz, hemisphere x trial two-way ANOVA, hemisphere effect, F_(1,6)_ = 0.98, p = 0.36, trial effect F_(19,114)_ = 1.17, p = 0.30, interaction effect F_(19,114)_ = 0.48, p = 0.97; lobule II/III-HPC: 9.08 ± 0.09 Hz, hemisphere x trial two-way ANOVA, hemisphere effect, F_(1,7)_ = 1.59, p = 0.25, trial effect, F_(19,133)_ = 1.33, p = 0.18, interaction effect, F _(19,133)_ = 0.97, p = 0.50; lobule VI-HPC: mean peak frequency = 8.78 ± 0.05 Hz, hemisphere x trial two-way ANOVA, hemisphere effect, F_(1,10)_ = 0.15 p = 0.70, trial effect, F _(19,190)_ = 1.21, p = 0.25, interaction effect, F _(19,190)_ = 0.53, p = 0.94). Further examination of the power and coherence spectra across a wider frequency range (1 to 45 Hz) in trials 1 and 20 confirmed that the observed changes across training were restricted to the theta band (Fig. 4 E-H).

To examine whether the observed changes in coherence across learning of the linear track were specifically related to performance of the goal-directed task itself, we next conducted pairwise analysis of cerebello-hippocampal theta coherence levels across the following conditions: home-cage prior to any linear track training (HC pre LT), first and last trials in the linear track (early and late LT), and home-cage following the end of training in the linear track task (HC post LT).

From the three cerebello-hippocampal recording configurations, only Crus I-HPC 6-12Hz coherence varied significantly across task conditions (Fig. 5, Crus I-HPC: Friedman test with FDR correction, Friedman statistic = 15.45, p = 0.0015). At the outset of linear track learning, HPC-Crus I coherence values did not significantly differ from home-cage (HC pre LT = 0.25 ± 0.02; early LT = 0.26 ± 0.02; HC pre LT vs early LT *q* = 0.44). However, during late stage linear track learning, the level of coherence was significantly higher than in home-cage recordings and early stages of learning (late LT = 0.29 ± 0.03; HC pre LT vs late LT *q* = 0.0015; early LT vs late LT *q* = 0.0016). When mice were returned to the home-cage environment following completion of linear track training (HC post LT) the level of HPC-Crus I coherence dropped significantly, back to pre-training levels (HC post LT = 0.26 ± 0.02; late LT vs HC post LT *q* = 0.012). We further analyzed changes in running speed and 6-12Hz power across conditions and found that their pattern of modulation was markedly different from the observed Crus I-HPC theta coherence dynamics (Fig. S8 and Fig. 5). While speed significantly varied between all conditions, 6-12 Hz power remained stable in both in the HPC and cerebellar recordings (Fig. S8). Together this suggests that the observed coherence dynamics appear to be at least partially independent of changes in speed or 6-12Hz oscillation power.

**Figure 5.**
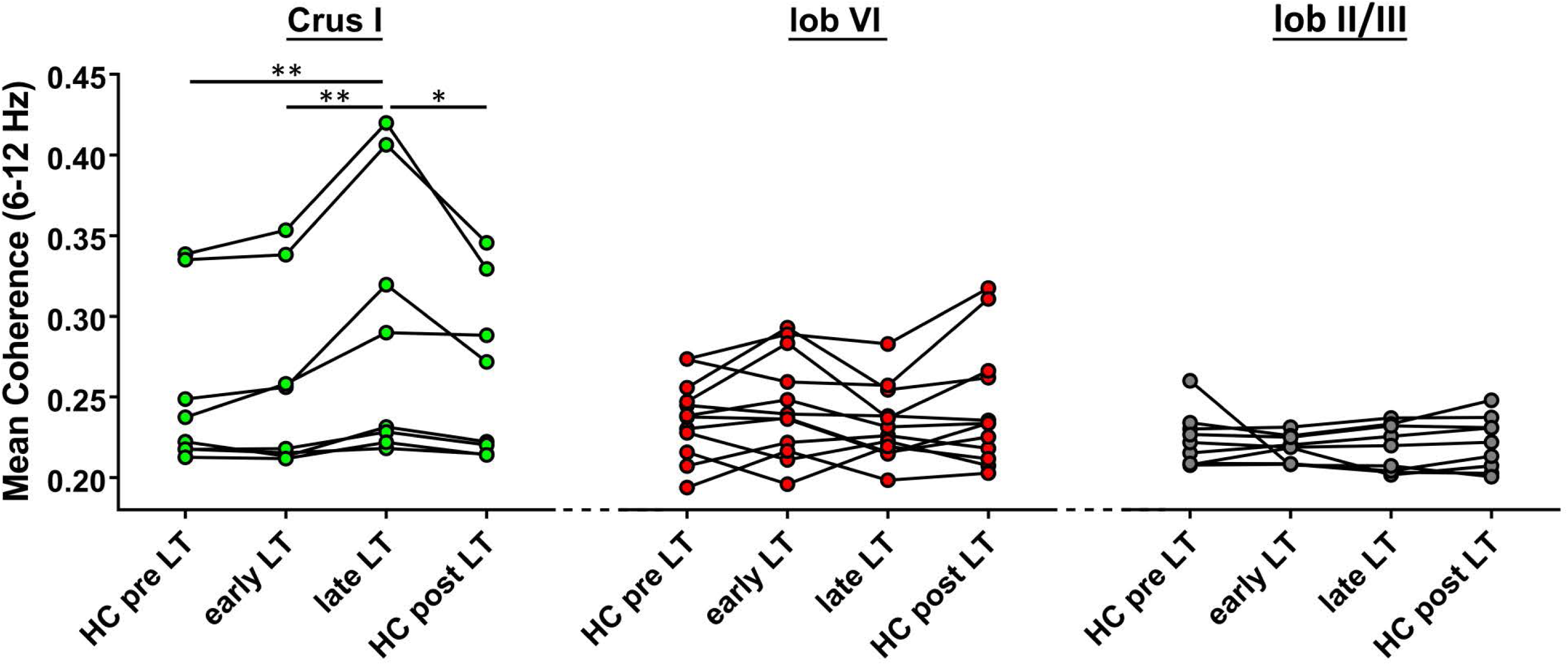
Cerebello-hippocampal interaction dynamics. Mean 6-12 Hz coherence for each cerebello-hippocampal recording combination (Crus I, n = 8 values/5 mice; lob II/III, n = 9 values/6 mice; lob VI, n = 12 values/7 mice) obtained during homecage recordings before training in the linear track (HC pre LT), during a representative trial of early training in the linear track (trial 1; early LT), during a representative trial of late training in the linear track (trial 20; late LT) and during homecage recording following completion of linear track training (HC post LT). A significant increase in Crus I–hippocampus coherence was observed during late LT compared to both HC pre LT and early LT, which then returned to pre-training levels during HC post LT (paired Friedman test with FDR correction, p = 0.0037; * q < 0.05, ** q < 0.01.). Lobule VI - hippocampus and lobule II/III - hippocampus coherence levels did not change significantly across conditions (paired Friedman test, FDR corrected p-value = 0.1718 and 0.5319, respectively). Abbreviations, lob II/III, lobule II/III; lob VI, lobule VI.

### Cerebello-hippocampal interactions during locomotion in a virtual environment

Our data indicate the presence of dynamic coherence between distinct cerebellar lobules and the dorsal hippocampus during goal-directed behavior. To investigate if this interaction requires the presence of specific sensory inputs, we further analyzed cerebello-hippocampal 6-12Hz coherence under conditions in which such inputs are not relevant for the behavioral task.

Head fixed mice were trained to locate rewards (medial forebrain bundle stimulation) at fixed positions on a virtual-reality based linear track (VR; Fig. 6A and B; see Methods). In this paradigm, vestibular, olfactory and whisker information cannot be reliably used to learn the task and thus it is likely that behavioral performance is linked mainly to visuo-motor information processing. Mice rapidly reached a stable performance level as illustrated by a stable running speed and number of rewards obtained (Fig. 6C) (mean speed: trial 4 = 7.61 ± 1.50 cm/s, trial 21 = 8.76 ± 2.78 cm/s, one-way repeated measures ANOVA, F_(1.789,8.945)_ = 0.82, *p* = 0.82; mean number of rewards: trial 4 = 25 ± 5, trial 21 = 36 ± 16, one-way repeated measures ANOVA, F_(1.519,7.596)_ = 0.93, *p* = 0.41; N = 6 mice). Although HPC theta peak frequency was variable across trials with stable behavioral performance (trials 4 to 21, Fig. S9A), mean 6-12Hz power and coherence values were similar (Fig. S9B-C) and therefore collapsed across these trials for further analysis (Fig. 6D-F)

**Figure 6.**
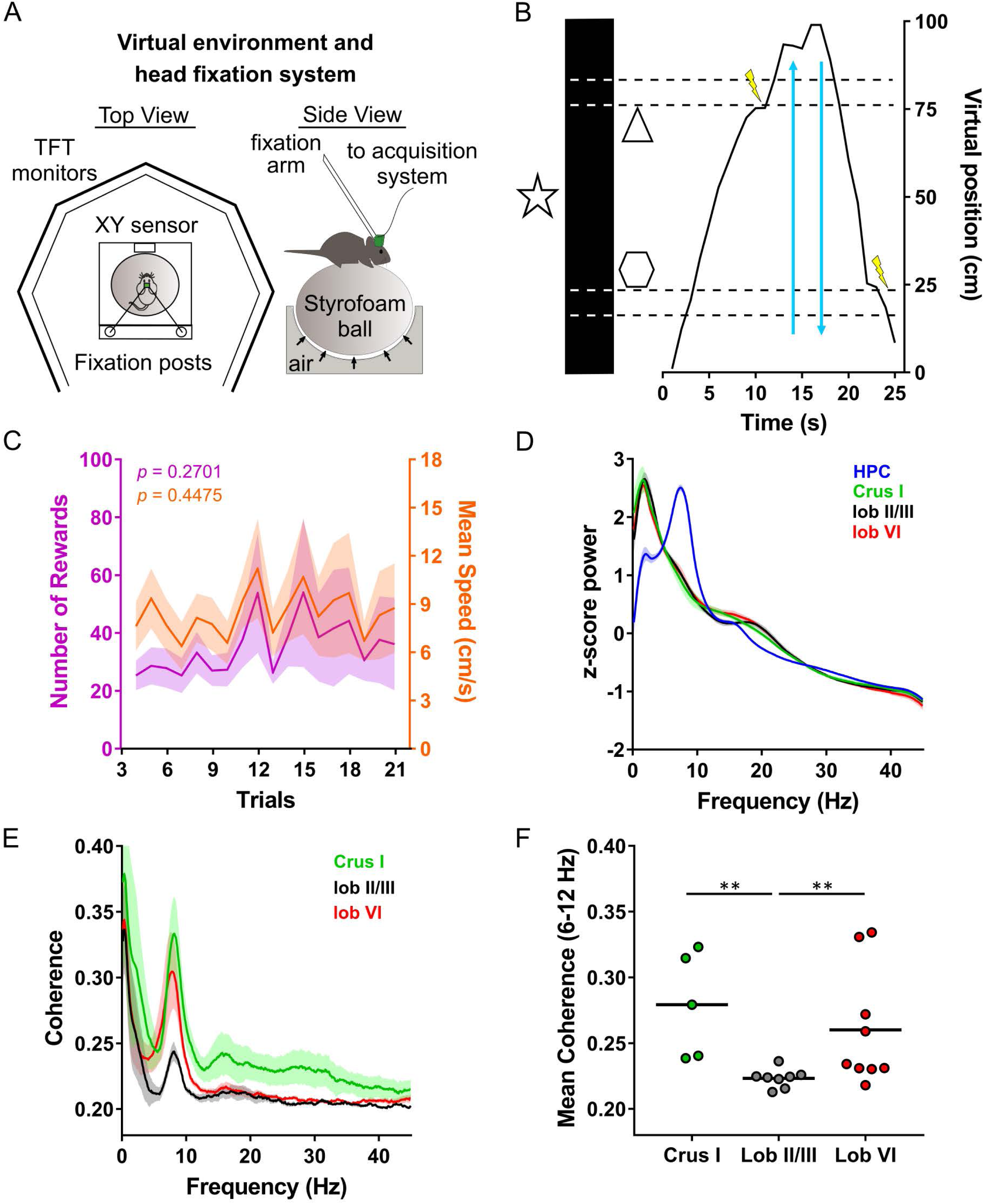
Cerebello-hippocampal coherence patterns during goal-directed behavior in virtual reality. **A**, Schematic of the virtual reality system and recording setup. Head-fixed mice were trained to move an air-cushioned Styrofoam ball in order to navigate through a virtual environment displayed on six TFT monitors surrounding the animal. **B**, Example recording of the virtual position as a mouse traversed a virtual linear track to receive rewards (MFB stimulation indicated by a lightning symbol, n = 7). **C**, Behavioural performance remained stable across trials as illustrated by the mean number of rewards (purple line; plotted against left Y axis; one-way repeated measures ANOVA, p = 0.4070) and the mean speed (orange line; plotted against the right Y axis; one-way repeated measures ANOVA, p = 0.4583). **D**, Pooled, z-scored normalized power spectra of hippocampal LFP recorded from the left (n = 16 mice) and right (n = 18 mice) hemispheres during homecage exploration and from cerebellar Crus I (n = 12), lobule II/III (n = 10) and lobule VI (n = 18). **E**, Averaged coherence between cerebellar cortical regions (colour coded) and bilateral hippocampus, pooled across all trials (Crus I-HPC, n = 5 values/3 mice; lobule II/III-HPC, n = 8 values/5 mice; lobule VI-HPC, n = 9 values/5 mice). **F**, Mean cerebello-hippocampal coherence in the 6-12 Hz frequency range. Both lobule VI and Crus I showed significantly higher coherence with hippocampus than lobule II/III (Kruskal Wallace with FDR correction, p = 0.0137; ** q < 0.01). Shading indicates S.E.M.

In keeping with results obtained in home-cage and real world (RW) linear track experiments, coherence spectra calculated between either Crus I, lobule VI or lobule II/III and HPC contained a single peak in the 6-12Hz theta frequency range (Fig. 6E). In this condition, 6-12Hz coherence levels between HPC and both Crus I and lobule VI were again significantly higher in comparison to lobule II/III (Fig. 6F; Kruskal-Wallis with FDR correction, H = 10.93, Crus I-HPC vs lobule II/III-HPC *q* = 0.0021, lobule VI vs lobule II/III *q* = 0.0077, Crus I vs lobule VI *q* = 0.2998; Crus I n = 5, lobule VI n = 9, lobule II/III n = 8).

In an effort to isolate the impact of changing sensory information on observed cerebello-hippocampal interactions from the influence of ongoing motor behavior, we next made pairwise comparisons of theta coherence values from recordings made during RW and VR linear track tasks in specific trials in which the number of rewards obtained and mean running speed was similar in both conditions (Fig. 7A; number of rewards: LT = 37 ± 4, VR = 36 ± 4, paired t test, t_5_ = 0.54, *p* = 0.46; speed: LT = 8.36 ± 0.47 cm/s, VR = 8.50 ± 0.56 cm/s, paired t test t_5_ = 0.68, *p* = 0.67; N = 6 mice). Patterns of coherence in the selected RW trials resembled those observed during late training in the LT (Fig. 7B; differences in mean 6-12 Hz coherence: one way ANOVA with FDR correction F_(2,19)_ = 4.55, Crus I-HPC vs lobule II/III *q* = 0.015, Crus I-HPC vs lobule VI-HPC *q* = 0.048, lobule VI-HPC vs lobule II/III-HPC *q* = 0.21; compare with Fig. 4H) and the selected VR trials closely mirrored the overall, pooled data (Fig. 7C; Kruskal-Wallis with FDR correction, H = 9.02, Crus I-HPC vs lobule II/III *q* = 0.009, Crus I-HPC vs lobule VI-HPC *q* = 0.33, lobule VI-HPC vs lobule II/III-HPC *q* = 0.007; compare with Fig. 6E), respectively. In these epochs of comparable motor state, Crus I-HPC coherence was reduced significantly in the VR condition compared to RW (Fig. 7D; RW = 0.2974 ± 0.030, VR = 0.2721 ± 0.022; paired t test T_4_ = 2.82, *p* = 0.047; n = 5). In contrast, lobule VI and lobule II/III – hippocampal coherence was similar across conditions (Fig. 7D; lobule VI: RW = 0.250 ± 0.011, VR = 0.260 ± 0.014, paired t test T_7_ =1.63, *p* = 0.39, n = 9; lobule II/III: RW = 0.239 ± 0.008, VR = 0.216 ± 0.002, paired t test, T_8_=0.86 *p* = 0.18, n = 8).

**Figure 7.**
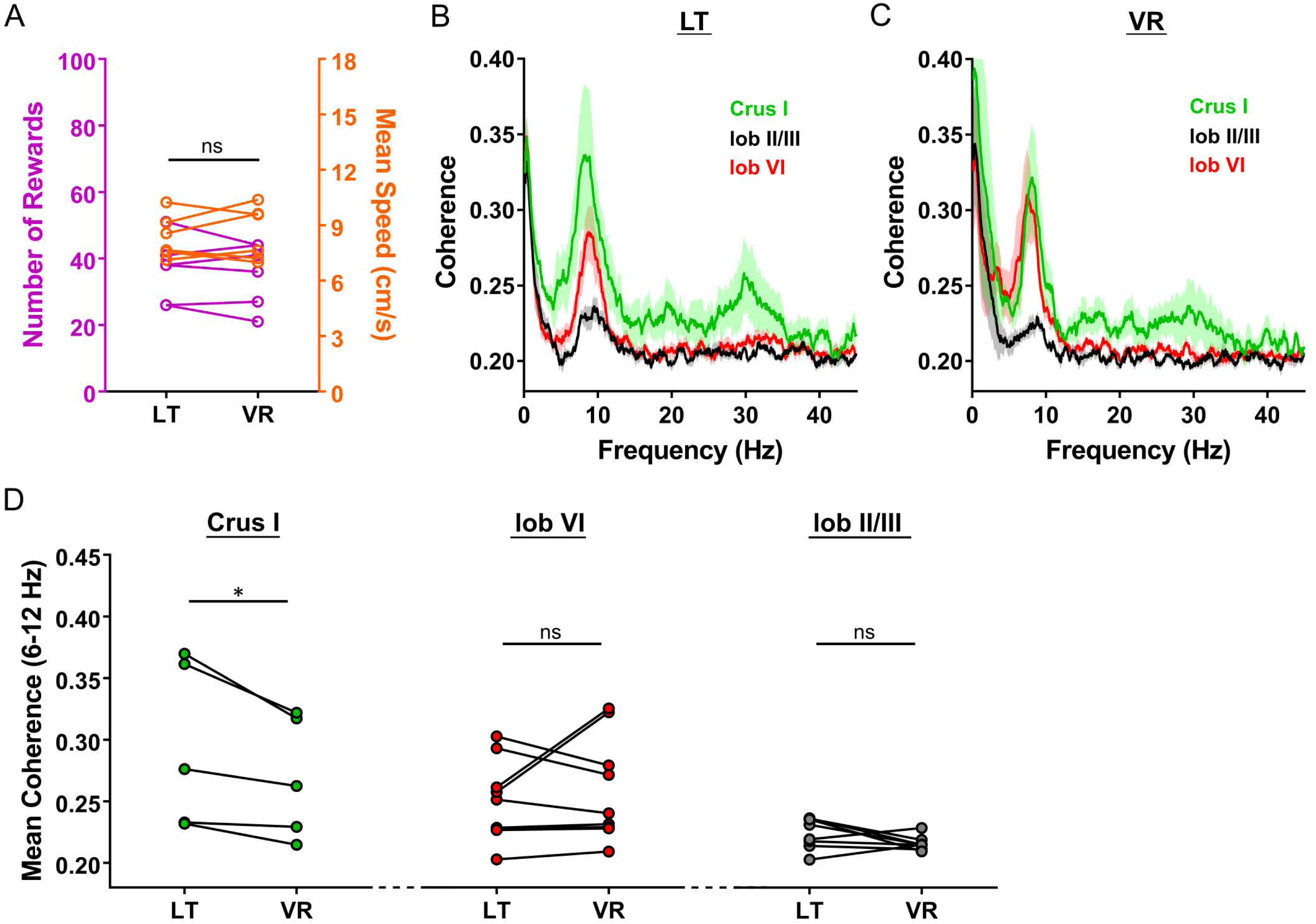
Comparison of cerebello-hippocampal interactions during epochs of similar behavioural performances in real-world and virtual-reality conditions. **A**, For each mouse (n = 6) we selected one trial from the real world linear track (LT) and one from the virtual reality (VR) condition in which behavioural performance was similar, as defined by non-significant changes in the number of rewards obtained (purple circles; plotted against left Y axis; paired t test, p = 0.5279) or the mean speed (orange circles; plotted against right Y axis; paired t test, p = 0.6119). **B**, Overall coherence between cerebellar cortical regions (colour coded) and bilateral hippocampus (Crus I, n = 5 values/3 mice; lobule II, n = 8 values/5 mice; lobule VI, n= 9 values/5 mice) in the selected trials from the linear track. **C**, Same as in **B** but for the selected trials from the virtual reality. **D**, For each recording combination, we compared 6-12Hz coherence values obtained from the selected linear track and virtual reality epochs. Crus I levels were significantly reduced in the virtual reality trials compared with the linear track (paired t test, *, p = 0.0480) while lobule VI or lobule II/III levels were similar in both conditions (Paired t tests, p = 0.1465 and 0.4165, respectively).

## Discussion

Taken together, our findings reveal previously undescribed cerebellar inputs to the hippocampus and offer novel physiological insights into a long-range neural network linking disparate brain regions initially assumed to support divergent behavioral functions, namely spatial navigation (hippocampus) and motor control (cerebellum). Topographically restricted regions of cerebellar cortex discretely route through restricted parts of their associated nuclei en-route to the hippocampus. Congruently, our physiological data reveal that these connected cerebellar regions dynamically interact with the hippocampus during behavior, via 6-12 Hz LFP coherence. Our findings thus offer an anatomical and physiological framework for cerebello-hippocampal interactions that could support cerebellar contributions to hippocampal processes [3], including spatial map maintenance [17,23,24].

Whilst previous studies provide compelling physiological evidence of cerebellar influences on the hippocampus [19–22,25,46], they do not provide the spatial resolution afforded by neuroanatomical tracing. Indeed, to the best of our knowledge, anatomical tracing studies have failed to report a mono-synaptic ascending cerebello-hippocampal projection. This is consistent with our rabies virus tracing study, in which incubation periods of 48-58 h were required before cell labeling was seen in the cerebellar nuclei. Such a timescale is indicative of a multi-synaptic pathway [35,47–49] potentially involving indirect connectivity through the forebrain navigation circuits.

Our anatomical results highlight three main inputs to the hippocampus emanating from the cerebellum. The first input we reveal originates from the vestibulo-cerebellum, specifically from the dorsal and ventral paraflocculus, which is likely routed via the vestibular and dentate nuclei [44]. This anatomical connection between the vestibulo-cerebellum and the hippocampus reinforces the already well described influence of the vestibular system on hippocampal dependent functions [50–52].

In addition to the classically described vestibular pathway, our data reveal that the central cerebellum also provides inputs to the hippocampus from vermal lobule VI, routed through caudal fastigial nucleus, and from Crus I, routed through the dentate. Using a combination of RABV expression and zebrin II staining we identified three specific cerebellar modules involved in these inputs: (1) the A module in lobule VI, (2) the hemispheric Crus I D2 module and (3) the Crus I paravermal C2 module. Of the latter two modules, C2 is likely less prominently anatomically connected with the hippocampus since the number of RABV+ cells in the nucleus interpositus posterior, its output nucleus [32], was minor compared with the other cerebellar nuclei. The convergence of inputs from disparate cerebellar zones (flocculo-nodular and central zones) and modules from vermal (A), paravermal (C2) and hemispheric (D2) regions in to the hippocampus suggest that its optimal function requires the integration of multiple aspects of sensory-motor processing carried out at these distinct cerebellar locations.

The A module in lobule VI is part of the so-called oculomotor vermis. It receives climbing fibers from the caudal medial accessory olive, and sends mainly ascending projections through the caudal portion of fastigial nucleus [53]. The oculomotor vermis receives multiple sensory inputs which include visual, proprioceptive, vibrissae, vestibular and auditory inputs conveyed by both climbing and mossy fibers [44]. It projects to, amongst others, the superior colliculus and other visual structures of the midbrain, the vestibular nuclei, the periaqueductal grey and the ventro-medial nucleus of the thalamus [54]. Notably, all of these regions contained RABV+ cells at 48h p.i., and thus they cannot be excluded as potential routes towards the hippocampus. The D2 module receives its climbing fiber input from the dorsal cap of the principal olive and projects out of the cerebellum through the rostromedial dentate nucleus [55]. It receives mossy fiber inputs carrying somatosensory, motor [56], and visual [57] information; along with inputs from the prefrontal cortex [7]. Climbing fiber inputs to this module relay information from the parvocellular red nucleus, which receives projections from premotor, motor, supplementary motor and posterior parietal areas. The majority of these cortical areas also receive projections from the D2 module after a thalamic relay in the ventro-lateral nucleus [7,58].

Complementary to these anatomical results, our electrophysiological findings reveal coherent activity between the hippocampus and those cerebellar lobules that are anatomically connected with it (lobule 6 and Crus I). This interaction was restricted to the 6-12 Hz range in the awake, behaving animal and showed non-uniform, dynamic profiles that were lobule dependent. Oscillations can align neuronal activity within and across brain regions, facilitating cross-structure interactions [e.g 33,59]. Cerebellar circuits support oscillations across a range of frequencies [for review see 60,61]. Of particular relevance to the current study are reports of oscillations within the theta frequency (∼4-12 Hz), which have been described in the cerebellar input layers at the Golgi [62] and granule cell [e.g. 63,64] level, and also in the cerebellar output nuclei [9,65]. Indeed, despite of the absence of prominent sustained theta band activity in our overall cerebellar recordings, we could record transient 6-12 Hz cerebellar oscillations (Fig S6).

Neuronal coherence has been described across the cerebro-cerebellar system at a variety of low frequencies [9,66–71] and oscillations within the theta range are thought to support inter-region communication across a wide variety of brain regions [72]. Our finding that cerebello-hippocampal coherence is limited to the 6-12 Hz bandwidth is in keeping with previous studies on cerebro-cerebellar communication in which neuronal synchronization has been observed between the cerebellum and prefrontal cortex [9,67], primary motor cortex [66,70], supplementary motor area [66] and sensory cortex [66]. Furthermore, local field potentials recorded in the hippocampus and cerebellar cortex are synchronized within the theta bandwidth during trace eye-blink conditioning in rabbits [73]. Human brain imaging studies have also described co-activation of blood oxygen level dependent signals in both cerebellar and hippocampal regions during navigation [15] and spatio-temporal prediction tasks [74], thus highlighting neuronal putative interactions between the two structures. Regarding studies in mice, a recent study has demonstrated the existence of statistically significant co-activation of the dorsal hippocampus and cerebellar lobules IV-V, lobule VI and Crus I after the acquisition of a sequence-based navigation task [16].

Multiple lines of evidence suggest that the phase-locking described here is unlikely to have resulted from volume conduction: 1) Rather than using a common reference electrode, our recordings were bipolar, with each recording electrode being locally and independently referenced [75]. 2) If volume conduction of theta oscillations was emanating from a hippocampal source then it could be assumed that cerebellar regions in closer proximity to the hippocampus would show higher levels of coherence (Fig. S5A). However, we found that coherence values were not related to the relative distance between the hippocampus and cerebellar recording site. 3) Hippocampal theta power increased over training in our linear track paradigm, whereas 6-12 Hz power remained stable across all cerebellar recording sites. However, significant coherence was only observed between hippocampus and Crus I. Thus, the observed coherence was unlikely to have resulted from co-variation in theta power between the two areas.

Importantly, we have shown for the first time that theta rhythms in the hippocampus preferentially phase lock with those in discrete regions of the cerebellum and that degree of this coupling changes depending upon the behavioral context. Lobule VI-hippocampus coherence was dominant during active movement in the home-cage and remained stable during learning of the real world linear track task. On the other hand, Crus I-HPC coherence was highly dynamic, showing a significant increase over the learning of the real world linear track task and becoming dominant after the acquisition of a goal-directed behavior. Interestingly, although multiple streams of sensory input, including those of a vestibular, whisker and olfactory nature, become irrelevant and even confounding in the head-restricted virtual environment task, Crus I-HPC and lobule VI-HPC coherence remained high in this condition. Paired-comparisons of trial epochs containing similar behavioural performances in the real-world and virtual environment revealed that Crus I-HPC coherence is significantly reduced in the latter while no change was observed for lobule VI-HPC.

We next consider our results within the modular understanding of cerebellar function. Within lobule VI, the A module receives multi-modal sensory information, mainly arising from collicular and vestibular centres [44]. The superior colliculus plays a role in visual processing and generation of orienting behaviors [76], which might be relevant for the establishment and maintenance of the hippocampal spatial map, and thus may be required constantly during active movement, independent of the specific behavioral task. The persistent and similar levels of lobule VI-HPC coherence during active movement in the homecage and linear track task, in both real world and virtual reality environment tasks is in agreement with such a hypothesis.

In monkeys and humans, Crus I is anatomically and functionally associated with prefrontal cortex [7,15]. In rat Crus I, the D2 module receives convergent sensory and motor information [77]. Furthermore, this module has been found to contain internal models, a neural representation of one’s body and the external world based on memory of previous experiences, that are used for visuo-motor coordination [78]. Similarly, the C2 module has been found to also participate in visuo-motor processing related to limb coordination during goal-directed reaching [79]. Both modules might be particularly important during the acquisition of a goal-directed behavior such as our real-world linear track task in which animals needed to reach non-cued reward zones. Our finding that Crus I-HPC coherence increases during task learning fits with this hypothesis. Furthermore, the observed reduction of Crus I-HPC coherence levels in the head-restricted, virtual environment task may reflect the reduced recruitment of cerebellar modules that are involved in processing of non-relevant sensory modalities, since only visuo-motor information can be reliably used to learn the task.

In summary, our results suggest the existence of both, an anatomically discrete hippocampal-cerebellar network interactions and a topographical dynamic weighting of these interactions potentially tailored to the prevailing sensory context and behavioral demands.

## Methods

Anatomical tracing studies were performed under protocol N°00895.01, in agreement with the Ministère de l’Enseignement Supérieur et de la Recherche. RABV injections were performed by vaccinated personnel in a biosafety containment level 2 laboratory.

All behavioral experiments were performed in accordance with the official European guidelines for the care and use of laboratory animals (86/609/EEC) and in accordance with the Policies of the French Committee of Ethics (Decrees n° 87–848 and n° 2001–464). The animal housing facility of the laboratory where experiments were made is fully accredited by the French Direction of Veterinary Services (B-75-05-24, 18 May 2010). Surgeries and experiments were authorized by the French Direction of Veterinary Services (authorization number: 75-752).

A total of 39 adult, male mice were used for this study. 18 adult male C57BL6-J mice were used for the anatomical tracing study, (Charles River, France) and 21 for the electrophysiology study (Janvier, France). 3 adult male CD-L7ChR2 mice were used for the dual hippocampal LFP and cerebellar unit-recording study (in-house colony derived from Jackson labs stock, USA).

Mice received food and water *ad libitum*, were housed individually (08: 00–20: 00 light cycle) following surgery and given a minimum of 5 days post-surgery recovery before experiments commenced.

### 1. Anatomy

#### Rabies virus injections

All the RABV (the French subtype of Challenge Virus Standard; CVS-N2C) inoculations were performed in the Plasticity and Physio-Pathology of Rhythmic Motor Networks (P3M) laboratory, Timone Neuroscience Institute, Marseille, France. Mice (n= 18) were injected intraperitoneally with an anesthetic mixture of ketamine (65 mg/kg; Imalgene, France) and xylazine (12 mg/kg; Rompun, Bayer) to achieve surgical levels of anesthesia, as evidenced by the absence of limb withdrawal and corneal reflexes and lack of whisking and were then placed in a stereotaxic frame (David Kopf Instruments, USA). The scalp was then incised, the skull exposed and a craniotomy drilled above the hippocampus.

Mice were injected with 200 nL of a mixture of one part 1% CTb Alexa Fluor^®^ 488 Conjugate (Invitrogen, distributed by Life Technologies, Saint Aubain, France) and four parts RABV in the left hippocampus (AP −2.0, ML +2.0, DV 1.97; Fig. 1, Fig. S1). Injections (200 nL/min) were performed using a pipette connected to a 10 μL Hamilton syringe mounted on a microdrive pump. Following infusion, the pipette was left in place for 5 min. The incision was then sutured and the animals allowed to recover in their individual home cage for either 30h (n= 5); 48 h (n= 3), 58h (n= 5) or 66 h (n=5). All animals were carefully monitored during the survival period and, in line with previous studies using these survival times, were found to be asymptomatic [48].

#### Tissue preparation

At the end of the survival time, mice were deeply anesthetized with sodium pentobarbitone (100mg/kg, intraperitoneal) then transcardially perfused with 0.9 % saline solution (15mL/min) followed by 75 mL of 0.1M phosphate buffer (PB) containing 4 % paraformaldehyde (PFA; pH = 7.4). The brain was then removed, post fixed for 2-3 days in 4 % PFA and then stored at 4°C in 0.1 M PB with 0.02% sodium azide. Extracted brains were then embedded in 3 % agarose before being coronally sectioned (40 μm) on a vibratome. Serial sections were collected and divided in 4 vials containing 0.1 M PB so consecutive slices in each vial were spaced 160 μm.

#### Injection site visualization

Sections from vial 1 were used to visualize the injection site by the presence of CTb. In most of the cases, the injected CTb was fluorescent and sections were directly mounted with Dapi Fluoromount G (SouthernBiotech^®^, Alabama, USA). In the other cases (S4-5, S11-13 and S17-18), the sections were first rinsed with PB 0.1M and then permeated with PB 0.1 M and 0.3 % Triton X-100. They were then incubated overnight in a choleragenoid antibody raised in goat (goat anti-CTb, lot no. 703, List Biological Laboratories, USA) diluted 1: 2000 in a blocking solution (PB 0.1 M, 5 % BSA). Subsequently, the sections were rinsed in PB 0.1M and incubated 4 h at room temperature with donkey anti-goat secondary antibody (1: 1000 in the blocking solution; Alexa Fluor^®^ 555, Invitrogen, distributed by ThermoFisher Scientific, Massachusetts, USA). Finally, they were also mounted with Dapi Fluoromount G.

The injection site was then visualized using a fluorescence microscope equipped with a fluorescein isothiocyanate filter (Axio Zoom V16, Carl Zeiss, France).

#### Rabies virus labeled cell quantification

Sections from vial 2 were used for quantification and 3 D reconstruction of the RABV labeled cells. Sections mounted on gelatin-coated SuperFrost ^®^Plus slides (Menzel-Glaser, Braunschweig, Germany) were first rinsed with PB 0.1 M and pre-treated with 3 % H_2_O_2_ during 30 minutes for blocking reaction against endogenous peroxidase. Following pretreatment, the sections were incubated overnight at room temperature with an anti-rabies phosphoprotein mouse monoclonal antibody [80] diluted at 1: 10000 in a blocking solution (PB 0.1 M, 0.1 % BSA, goat serum 2 % and 0.2 % Triton X-100). The day after, the sections were rinsed in PB 0.1M and incubated 2 hours with a biotinylated affinity-purified goat anti-mouse IgG (1: 2000 in blocking solution; Santa-Cruz, Heidelberg, Germany). Then, they were also incubated using an avidin-biotin complex method (Vectastain Elite ABC-Peroxidase kit R.T.U. Universal, Vector Laboratories, Burlingame, CA, USA) to enhance sensitivity. For visualization, the sections were incubated in a 3,3’-diaminobenzidine-tetrahydrochloride (DAB) solution (0.05 % DAB and 0.015 % H_2_O_2_ in PB 0.1 M). Finally, they were counterstained with Cresyl and cover-slipped.

Quantitative analyses of rabies-positive nuclei were performed using a computerized image processing system (Mercator, Exploranova, France) coupled to an optical microscope. The quantification of rabies-positive nuclei was carried out at 10x magnification. Structures were defined according to a standard atlas [81]. Immunoreactive neurons were counted bilaterally. Representative images were obtained using an Axio Zoom V16 microscope (Carl Zeiss, France).

#### 3-D reconstruction

A Nikon Eclipse E800 microscope equipped with a digital color camera (Optronics, USA) was used to visualize mounted cerebellar sections under brightfield illumination. The contour of every 4 th section was then manually drawn using Microfire software (Neurolucida, MBF Bioscience, USA) and cell counts were performed. The sections were then aligned and stacked (160 µm spacing).

#### Rabies virus-zebrin II double immuno-staining

For case S18, sections from vial 3 were mounted on gelatin-coated SuperFrost ^®^Plus slides (Menzel-Glaser, Braunschweig, Germany), rinsed with PB 0.1M and then permeated and blocked in a solution of PB 0.1 M, 0.2 % Triton X-100 and bovine serum 2.5% for 30 minutes. Then they were incubated during 48h at 4°C in a mix of rabbit polyclonal anti-Aldolase C primary antibody (a kind gift from Izumi Sugihara [41]; No. 69075; 1:500000) and the mouse anti-rabies antibody used for the single RABV staining (1:5000) in a blocking solution (PB 0.1 M, 0.1 % Triton X-100 and bovine serum 1 %). Subsequently, the sections were first rinsed with PB 0.1 % and then incubated in a mix of RRX-Goat anti-rabbit IgG (1: 5000; ref 111-295-144, Jackson Immuno Research) and donkey anti-mouse secondary antibody (1: 5000; Alexa Fluor^®^ 647, Invitrogen distributed by ThermoFisher Scientific, Massachusetts, USA) in blocking solution. Finally, they were also mounted with Dapi Fluoromount G.

Images were obtained using an Axiozoom v16 microscope (Carl Zeiss, France) and contour and labeled neurons in the cerebellum were manually draw for reconstruction of the zebrin bands and cerebellar modules and location of the RABV+ cells.

### 2. Electrophysiology procedures

#### Subjects and surgical protocols

Bipolar LFP recording electrodes (interpolar distance of ∼0.5mm; 140µm diameter Teflon coated stainless-steel, A-M system, USA) were stereotaxically targeted to hippocampus (AP - 2.2, ML +2.0, DV 1.0), lobule 6 (AP −6.72, ML 0.0, DV 0.1), lobule 2 (AP −5.52, ML 0.0, DV 1.8) and Crus I (AP −6.24, ML 2.5, DV 0.1) of 21 C57BL6-J mice. Pairs of flexible stainless-steel wires were used to also record neck EMG (Cooner wire, USA)

In 15 C57BL6-J mice, bipolar stimulation electrodes (140-μm-diameter stainless steel; A-M system, USA) were also implanted at the left medial forebrain bundle [MFB; to serve as a reward signal; AP −1.4, ML +1.2, DV +4.8 [Cf. 82,83]. All electrode assemblies were fixed to the skull using a combination of UV activated cement (SpeedCem, Henry Shein, UK), SuperBond (SunMedical, Japan) and dental cement (Simplex Rapid, Kemdent, UK). Four miniature screws (Antrin, USA) were also attached to the skull for additional support and to serve as recording ground.

In 6 mice, a lightweight metal head fixation device (0.1g) was also implanted. The total implant weight did not exceed 2.5g (including head fixation post and cement).

#### Recording

Signals from all electrodes were attached to an electronic interface board (EIB 18, Neuralynx, USA) either during surgery. Differential recordings were made via a unity-gain headstage preamplifier (HS-18; Neuralynx, USA) and Digital Lynx SX acquisition system (Neuralynx, USA). LFP and EMG Signals were bandpass-filtered between 0.1 and 600 Hz and sampled at 1 kHz. Mouse position was tracked at 30Hz using video tracker software and infra-red LEDs attached to the headstage (Neuralynx, USA).

#### MFB Stimulation

Intracranial rewarding stimulation consisted of a 140Hz stimulation train lasting 100ms delivered through the headstage to the implanted electrodes (SD9k, Grass Technologies, USA). Optimal voltage for intracranial MFB was determined for each mouse with a nose-poke task prior to training (range, 1-6V [Cf. 82]).

#### Histology

After completion of all the experiments, mice were deeply anesthetized with ketamine/xylazine solution (150mg/kg) and electrolytic lesions created by passing a positive current through the electrodes (30µA, 10sec). With the electrodes left *in situ*, the animals were perfused transcardially with saline followed by paraformaldehyde (4%).

Brains were extracted and post-fixed in paraformaldehyde (4%; 24h) then embedded in agarose (24h). A freezing vibratome was used to cut 50μm thick sagittal cerebellar and coronal hippocampal sections. The sections were mounted on gelatinised slides and stained with cresyl violet. Recording locations were identified by localised lesions in the cerebellum and hippocampus and plotted on standard maps with reference to a stereotaxic atlas [81].

### 3. Behavioral procedures

#### Familiar environment

All recordings were made in the animal’s home-cage (30 cm x 10 cm x 10 cm plastic box), with the lid removed and lasted a maximum of 4 hours. Recordings were made during the day between the hours of 10 am to 18:00 pm.

#### Linear track – real world

The linear track was made in-house from 100 cm x 4 cm x 0.5 cm of black plastic positioned 20cm above the surface of the experimental table. The behavioral assembly was located in a separate room from the experimenter and was surrounded on four sides by black curtains. Three salient visual cues were placed at fixed locations along the edge of the track (10 cm from the edge). Mice were trained to run in a sequential manner from one end of the track to the other in order to receive a reward, which consisted of an electrical stimulation of the MFB. The reward stimulation was delivered automatically when the mice reached a 5 cm wide goal-zone, which was located 10 cm from the end of the track. Timing of the reward signal was logged on the electrophysiological recordings via TTL signals. Sessions lasted 12 mins and were repeated 3 times per day with an inter-session time of 5 mins over 7 days. Between sessions, the track was cleaned with 20 percent ethanol.

#### Linear track – virtual reality environment

A commercially available virtual-reality environment was used (Jet Ball, Phenosys, Germany), utilising an air cushioned Styrofoam ball (200 mm), which served as a spherical treadmill for head restrained mice [Cf. 84] (Supplementary Fig. 12). The floating ball assembly was positioned 20 cm from a series of six octagonally arranged TFT surround monitors (19 inch) such that the head restrained mice had an unobstructed view of the visual scene. Rotation of the Styrofoam ball was detected by an optical sensor (sampling frequency 5700 dots per inch at 1 kHz). The vertical axis signals were interpreted by the VR software as the forward and backward movement of the virtual position of the animal. Position within the VR was then translated to a voltage signal (zero to five volts, with 5 volts indicating the end of the track), and sent to the Digital Lynx SX (Neuralynx, USA) electrophysiology system via a DACQ interface (DACQ 6501, National Instruments, USA). The start of the VR display was logged on the electrophysiology recordings via a TTL signal. To provide a reward signal, when the mice reached a given location within the VR (10 cm from the end of the track) a TTL marker was sent to both the electrophysiological recording system (to provide a timestamp-marker of the event) and an electrical stimulus generator linked to the HS-18 headstage (in the same manner as for RW linear track experiments).

The virtual scene consisted of a 1 m long track with grey walls and included 3 salient visual cues. After 3 x 12 mins sessions of habituation to the head fixation on the floating-ball assembly, mice were trained to run on the linear track in 12 min sessions, 3 times per day with an inter-session interval of approximately 5 mins during 7 days. The number of rewards received by the animal was logged in the electrophysiology software (Cheetah 5.6.3, Neuralynx, USA).

### Behavioral and electrophysiological analysis

All data were processed in Matlab (Mathsworks, USA), Spike 2 (Cambridge Electronic Design, UK) and Prism (Graphpad, USA).

#### 1. Behavior

In the home-cage environment, behavioral data were selected using a custom-made Matlab script. Interactive cursors were used to define periods of active movement, based upon speed (derived from video tracking data), EMG and LFP signals. For the purpose of further analysis, we focused on periods of active movement (indicated by high EMG amplitude, speed and hippocampal 6-12 Hz oscillations). The overall mean speed of each mouse was then calculated across all the selected data epochs.

For RW linear track experiments, in each 12 minute trial (3 trials per day) the number of rewards (indicated by TTL markers), distance traveled, speed and an efficiency score (distance traveled per reward) were calculated using a custom Matlab script. These parameters were calculated on a trial-by-trial basis (from trial 2 onwards).

In virtual reality-based experiments, for each 12min trial (3 trials per day), the number of rewards (timestamped by TTL markers) and virtual speed was calculated using a custom Matlab script. Virtual speed was calculated using the virtual environment X and Y coordinate values (recorded as voltage signals in Neuralynx Cheetah software).

#### 2. Electrophysiology

Multi-taper Fourier analyses (Chronux toolbox [85]) were used to calculate power and coherence of the LFP data. We used a 10s sliding window (9s overlap) and 19 tapers for all analysis, except for the example presented in Fig. S6 in which we used a 2s sliding window (1.5s overlap) and 3 tapers. Statistical comparison was restricted to the 6-12 Hz frequency band unless otherwise stated.

For recordings made in the home-cage environment, LFP data were manually selected, as described above for behavioral data. Analysis was focused on LFP gathered during periods of active movement. Selected data were then filtered to remove any large-amplitude, low frequency artifacts using a stationary wavelet transform [86]. Spectral power and coherence were then calculated. The spectral power between 0.1 and 45 Hz was z-scored due to inter-animal variations in LFP magnitude. Mean power and coherence were calculated in the 6-12Hz frequency range for all cerebello-hippocampal combinations. Data duration in the home-cage environment varied across mice (range, 12 to 132 min). Therefore, to reduce the impact of data length on subsequent analyses and also to match with subsequent linear track experiments (duration of 12 min), for each mouse we concatenated the LFP in to 12 min blocks. When multiple 12 min blocks were available (number of data blocks ranged from 1 to 11) we calculated the average coherence across all blocks. The number of 12 minute blocks used was found to have no correlation with the overall level of calculated coherence (See Fig. S5F).

For RW and virtual linear track recordings, general electrophysiological analysis methods were the same as described for home-cage recordings including artifact removal procedures (in this case MFB stimulation artifacts were also removed using the methodology described in [86]). In addition to pooled calculations (in which analysis was conducted across all trials of the task), power and coherence was also calculated on a trial-by-trial basis across learning.

### Statistical analysis

Statistical analyses were conducted using Matlab Statistical Toolbox and Prism (Graphpad, USA). Normality was assessed using a Shapiro-Wilk test. Parametric and non-parametric tests were then used accordingly.

## Acknowledgements

This work was supported by the Fondation pour la Recherche Médicale DEQ20160334907-France, by the National Agency for Research ANR-17-CE16-0019-03 (LRR), by the CNRS and Aix-Marseille Université through UMR 7289 (PC). This work also received support under the program Investissements d’Avenir launched by the French Government and implemented by the ANR, with the references, PER-SU (LRR) and ANR-10-LABX-BioPsy (LRR). The group of LRR is member of the Labex BioPsy and ENP Foundation. Labex are supported by French State funds managed by the ANR within the Investissements d’Avenir programme under reference ANR-11-IDEX-0004-02. We thank Roxanna Ureta for help with histology, Lilith Sommer for help with behavioral experiments, Gregory Sedes and Nadine Francis for help developing analysis codes, and Richard Apps for his insightful comments. We are grateful to Richard Hawkes and Izumi Sugihara for generously providing the aldolase C antibody. Finally, we thank all members of the CEZAME team for helpful discussions of the experiments and manuscript.

## Figure Legends

**Figure S1.**
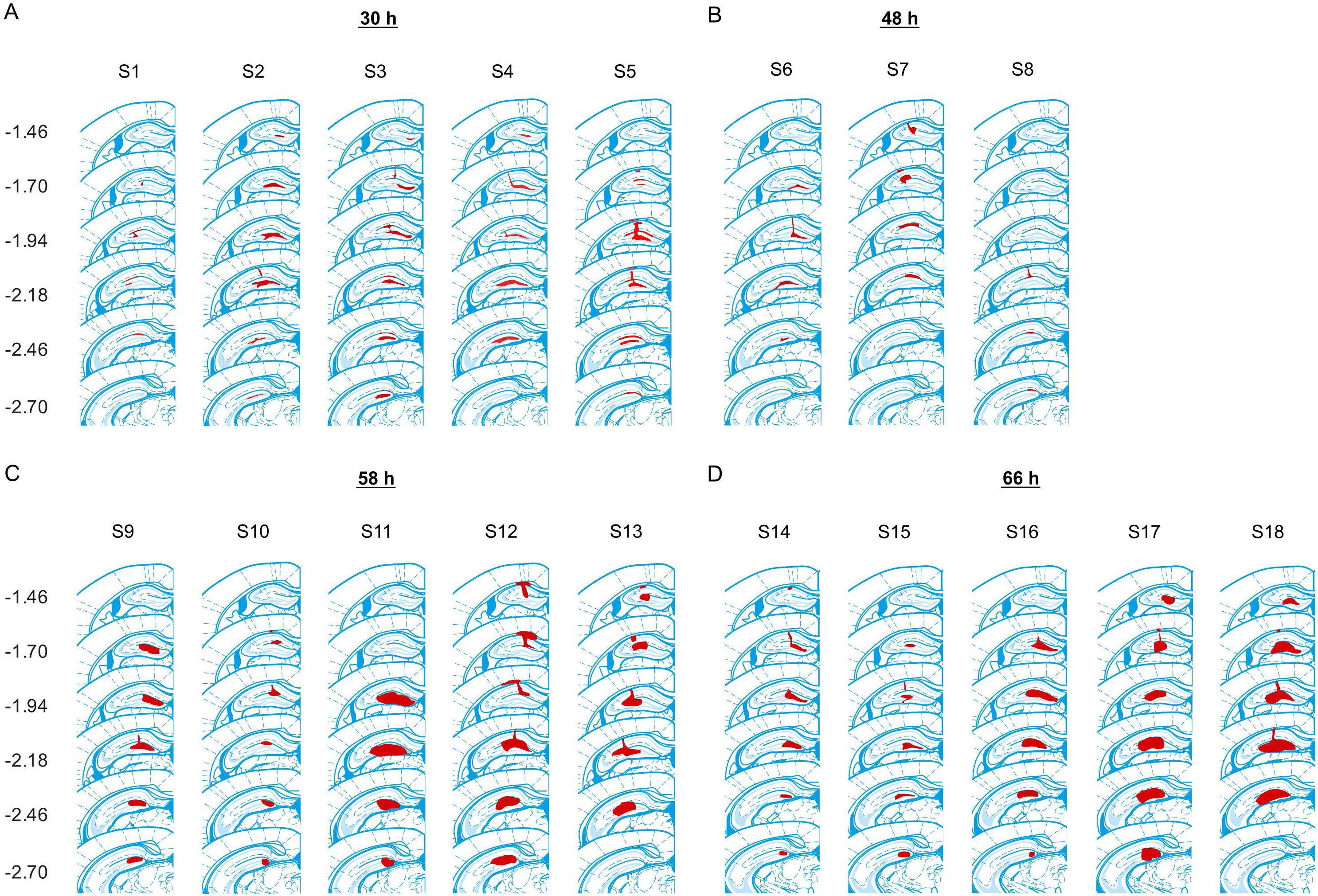
Location of rabies virus injection for the 4 experimental groups. RABV was co-injected with fluorescent CTB to visualize the injection spread. The location of the injection is indicated by the red area on a standard coronal outline of the left hippocampus adapted from [81]. Rostro-caudal position relative to bregma is indicated on the left (mm). Experimental ID for each case is shown above the sections. **A**, Injection sites of the 5 mice from the 30h survival time group **B**, Injection sites of the 3 mice from the 48h survival time group. **C**, Injection sites of the 5 mice from the 58h survival time group. **D**, Injection sites of the 5 mice from the 66h survival time group.

**Figure S2.**
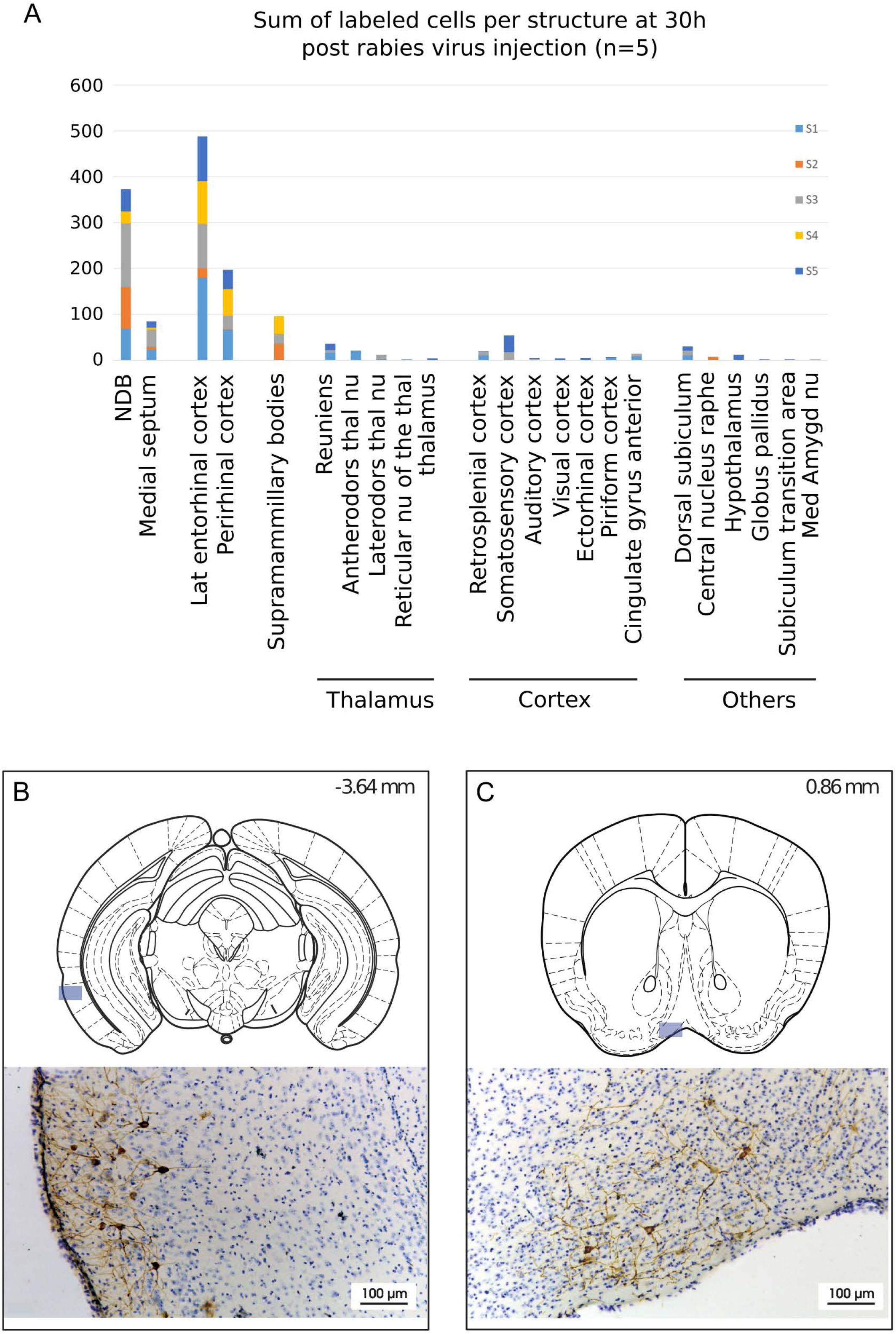
Rabies virus main labelled structures 30h after hippocampal rabies injection. **A**, Cumulative sum of labelled cells per structure after 30h post hippocampal rabies injection (colour coded for each case, n = 5 mice). **B-C,** Localisation and representative photomicrographs of RABV most labelled regions at 30h, lateral entorhinal cortex (**B**) and nucleus of the diagonal band (**C**). The position is indicated by a blue insert on a standard coronal brain section adapted from [81], and rostro-caudal position according to Bregma is indicated in the top-right corner. Abbreviations, ADN, antero-dorsal nucleus of the thalamus; LDN, latero-dorsal nucleus of the thalamus; Lat entorhinal cortex, lateral entorhinal cortex; NDB, nucleus of the diagonal band; TRN, thalamic reticular nucleus.

**Figure S3.**
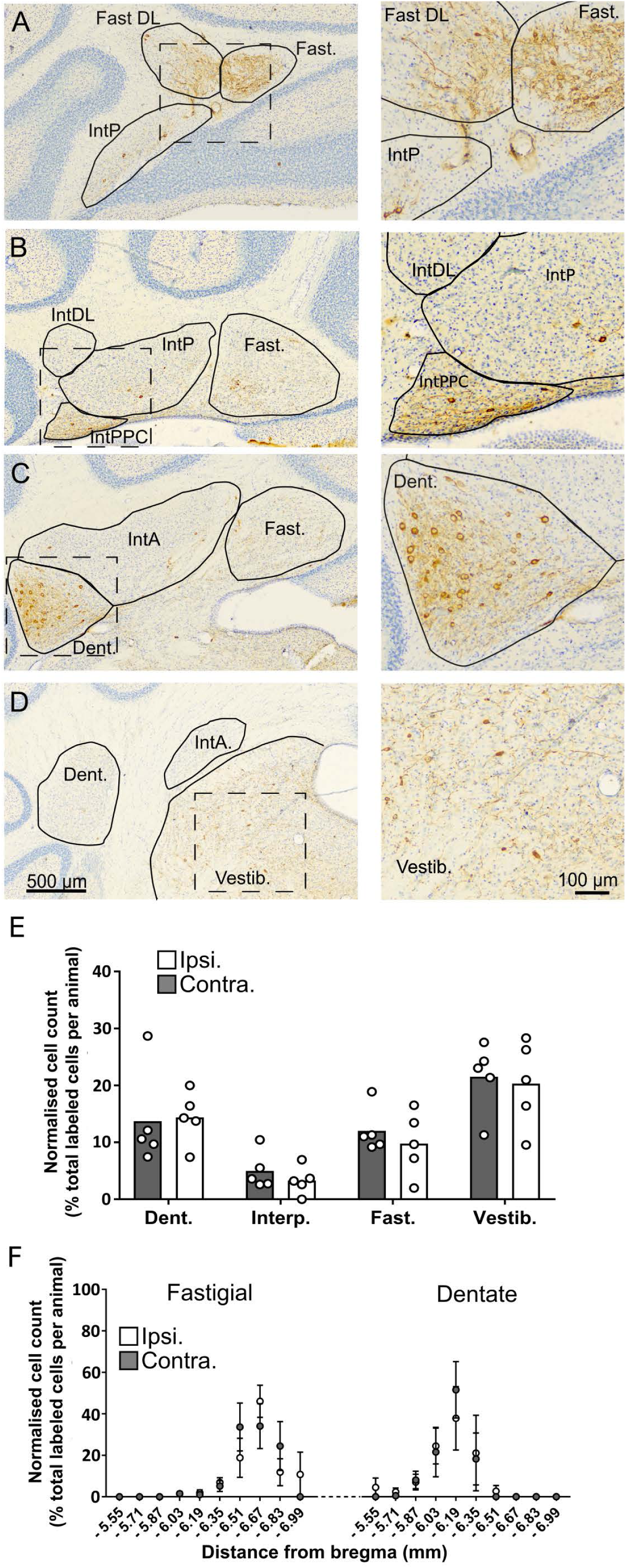
The topographical distribution of DCN labelling at 66h. **A-D**, representative photomicrographs showing labeling in the ipsilateral cerebellar and vestibular nuclei 66h after hippocampal rabies injection. Left panels show low magnification view, right panels show high magnification view of area indicated by dashed box. **E**, Pooled, normalized counts of rabies labeled cells in the ipsi-and contralateral cerebellar and vestibular nuclei (n = 5 mice). No significant differences were found between ipsi-and contralateral nuclei (nuclei x hemisphere two-way ANOVA, hemisphere effect F _(1,4)_ = 1.14, p = 0.35, interaction effect F _(3,12)_ = 0.21, p = 0.89, nuclei effect F _(3,12)_ = 7.88, p = 0.004). **F**, Normalized cell counts in the fastigial nucleus (left) and dentate nucleus (right) according to their rostro-caudal position relative to Bregma. Open circles, contralateral count; filled circles, ipsilateral count (n = 5 mice). Abbreviations: Dent., Dentate nucleus; Fast., fastigial nucleus; Fast. DL, dorsolateral fastigial nucleus; Interp., nucleus interpositus; IntA, nucleus interpositus anterior; IntDL, dorsolateral nucleus interpositus; IntP, nucleus interpositus posterior; Vestib., vestibular nuclei.

**Figure S4.**
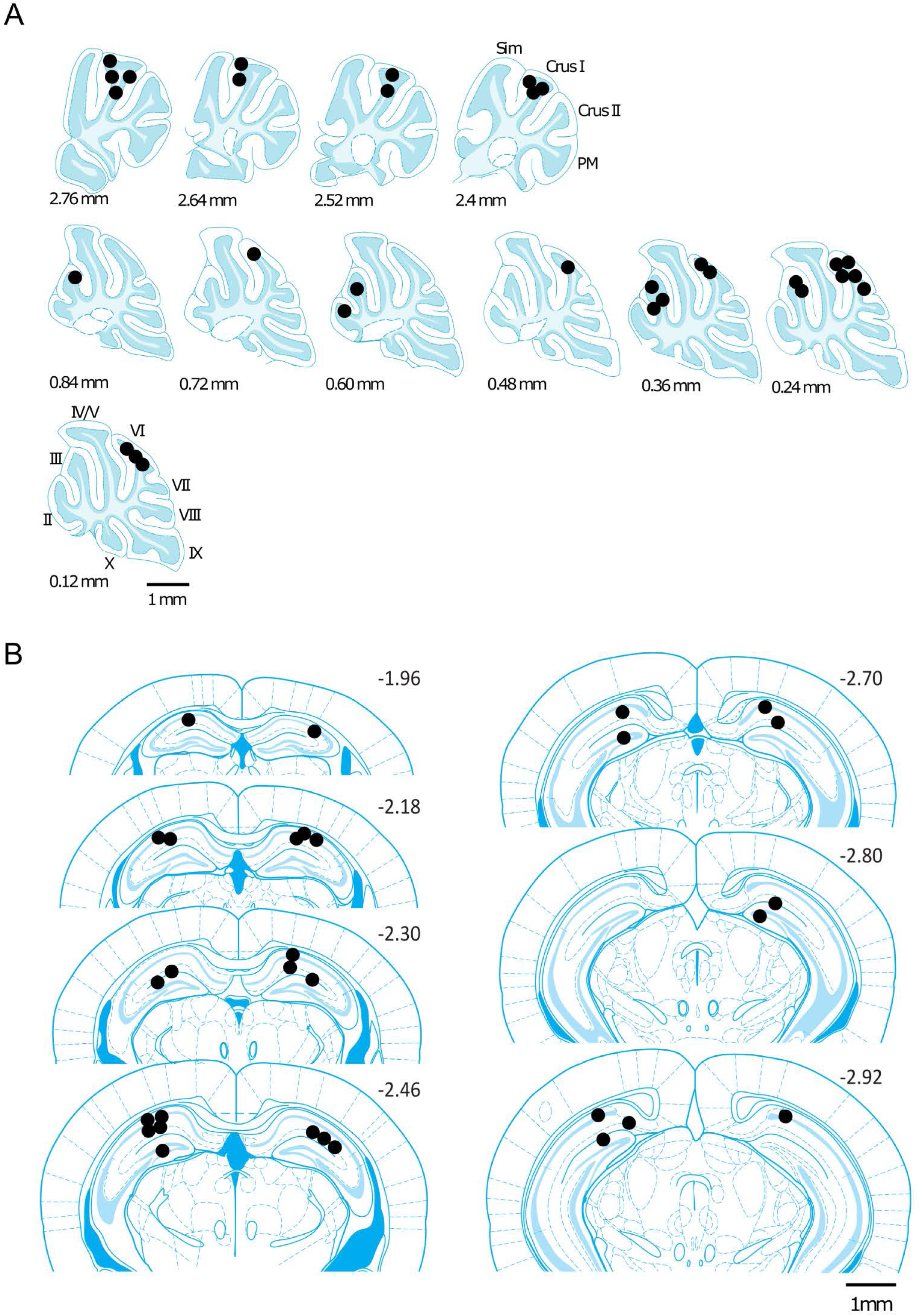
Reconstructed location of the implanted electrodes. The position of the implanted electrodes are represented by black dots on a standard sagittal outline of the cerebellum (**A**) or coronal outline of the hippocampus (**B**) [adapted from 81]. The medio-lateral (in **A**) or rostro-caudal (in **B**) positions relative to midline or bregma, respectively, are indicated next to the outlines. Abbreviations, II, lobule II; III, lobule III; IV/V, lobule IV/V; VI, lobule VI; VII, lobule VII; VIII, lobule VIII; IX, lobule IX; X, lobule X; Sim, lobule simplex; PM, paramedian lobule.

**Figure S5.**
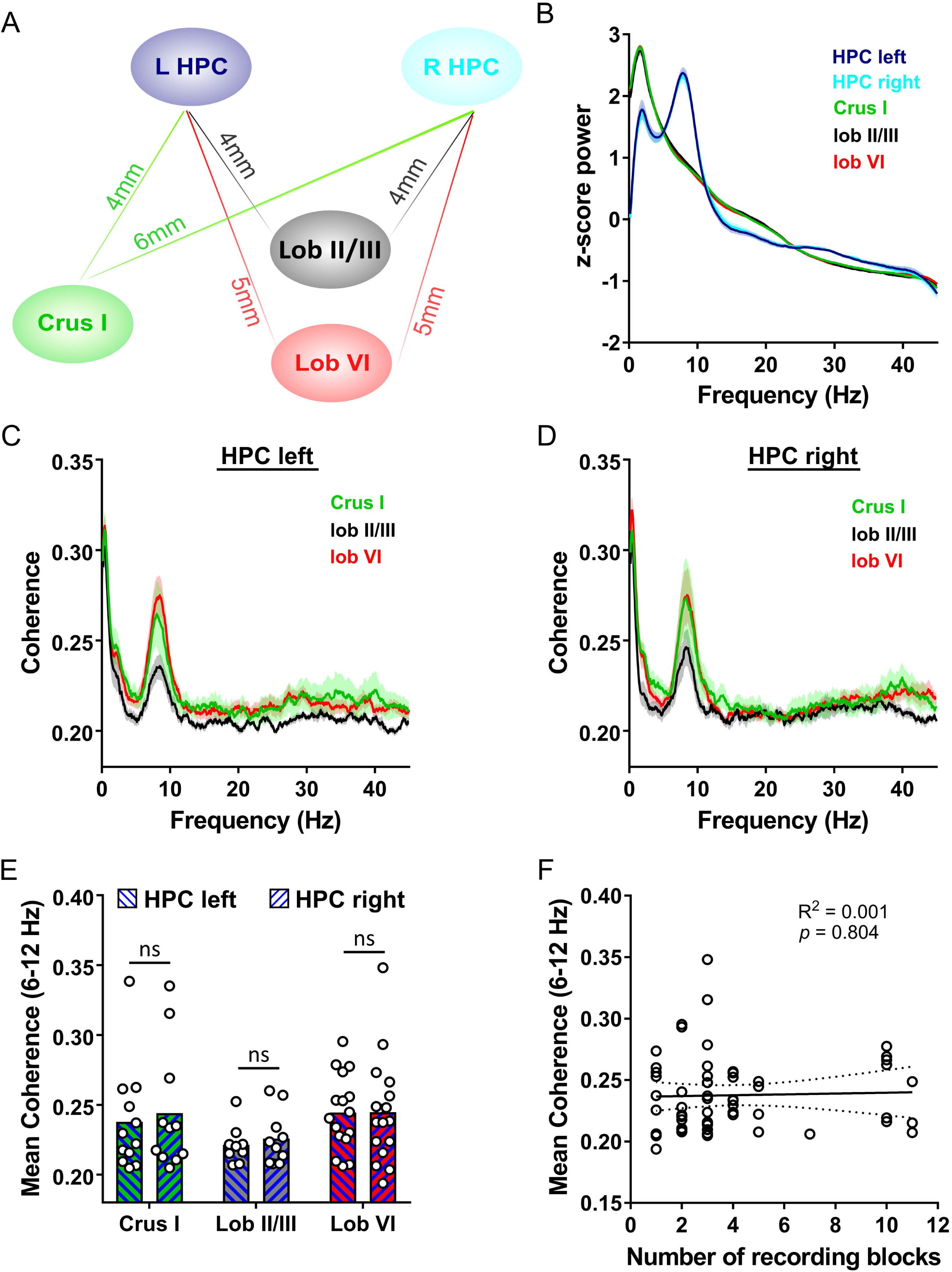
Cerebello-hippocampal coherence patterns are similar across hemispheres during active movement in homecage. **A**, Schematic indicating approximate distances between recording sites in the cerebellum and left/right dorsal hippocampus (HPC). **B**, Overall power spectra from right (n = 18) and left (n = 16) HPC and cerebellar recordings made from Crus I (n = 12), lobule II/III (n = 10) and lobule VI (n = 18) during active movement in the homecage environment. **C**, Overall coherence between cerebellar cortical regions (Crus I, n = 11; lobule II, n = 8; lobule VI, n = 15; colour coded) and left HPC during active homecage movement. **D**, Overall coherence between cerebellar cortical regions (Crus I, n = 10; lobule II, n = 9; lobule VI, n = 16; colour coded) and right HPC. **E**, Mean 6-12 Hz coherence between cerebellar recordings and left or right HPC. No differences were observed between hemispheres (hemisphere x combination one-way ANOVA with FDR correction, hemisphere effect p = 0.6355). **F**, The number of 12min analysis blocks was not correlated to the mean 6-12 Hz level of coherence obtained. Coherence values shown for all cerebello-hippocampal recording combinations (n= 57 values/23 mice; solid line indicates linear regression; dashed lines indicate 95% confidence intervals).ShadingindicatesS.E.M.

**Figure S6.**
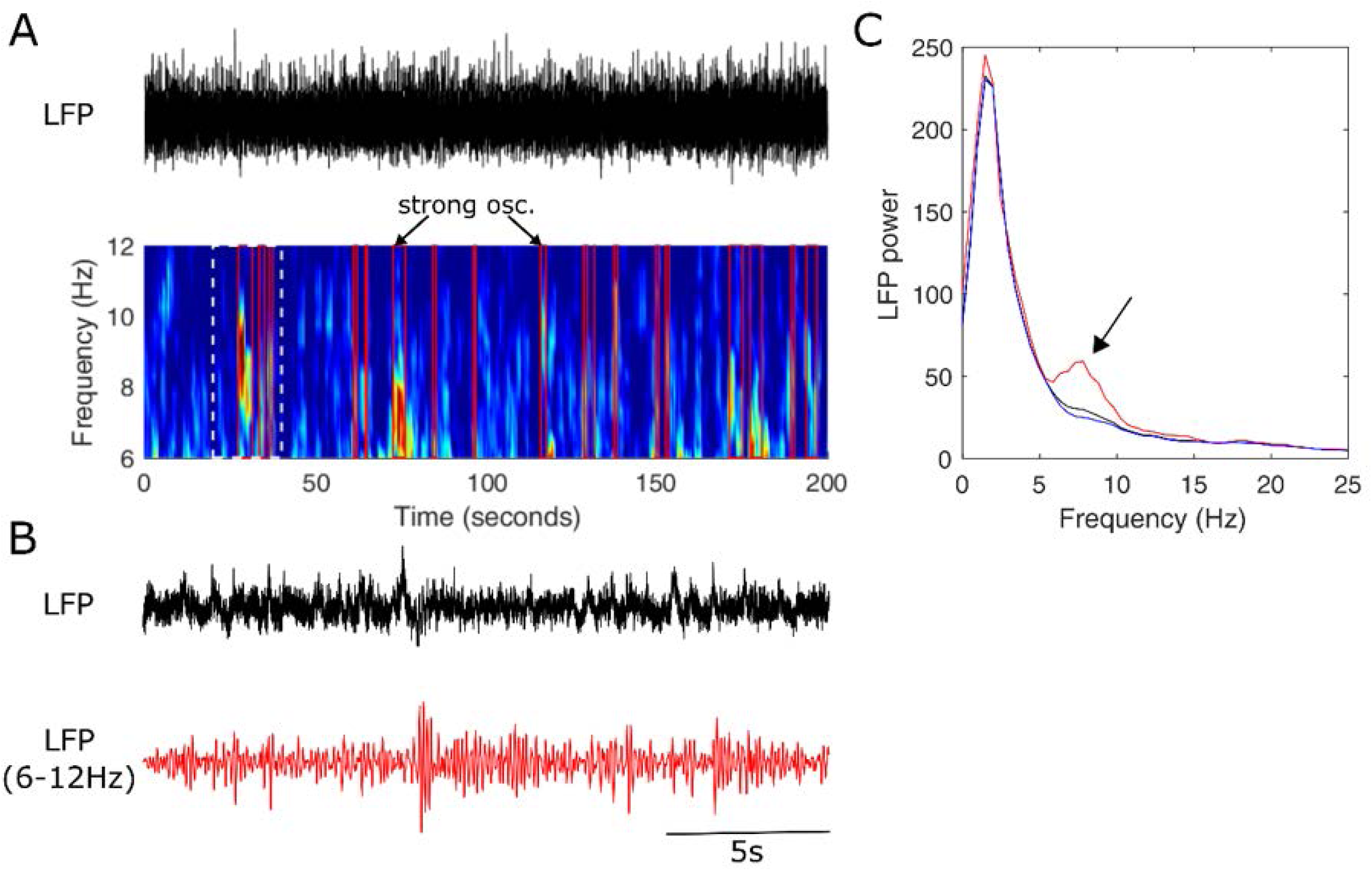
Transient 6-12 Hz oscillations are present in the cerebellar cortex. **A**, Upper trace, 200s epoch of LFP (0.1 to 600Hz) recorded from lobule VI during active movement in the homecage environment. Lower panel, spectrogram of the LFP. Periods of intense spectral power in the 6-12Hz band demarked by red boxes. **B**, 20s period of intense oscillation (from area marked by dashed white lines in **A**) is shown on a larger timescale. In red, same epoch filtered in the 6-12 Hz frequency band. **C**, Power spectra calculated from the whole period (black line), from epochs of high intensity in the 6-12Hz frequency band (epochs demarked by red boxes in A; red line) and from epochs without high intensity in the 6-12Hz frequency band (blue line). Arrow indicates peak in spectra within the 6-12Hz range, apparent in the selected high intensity periods.

**Figure S7.**
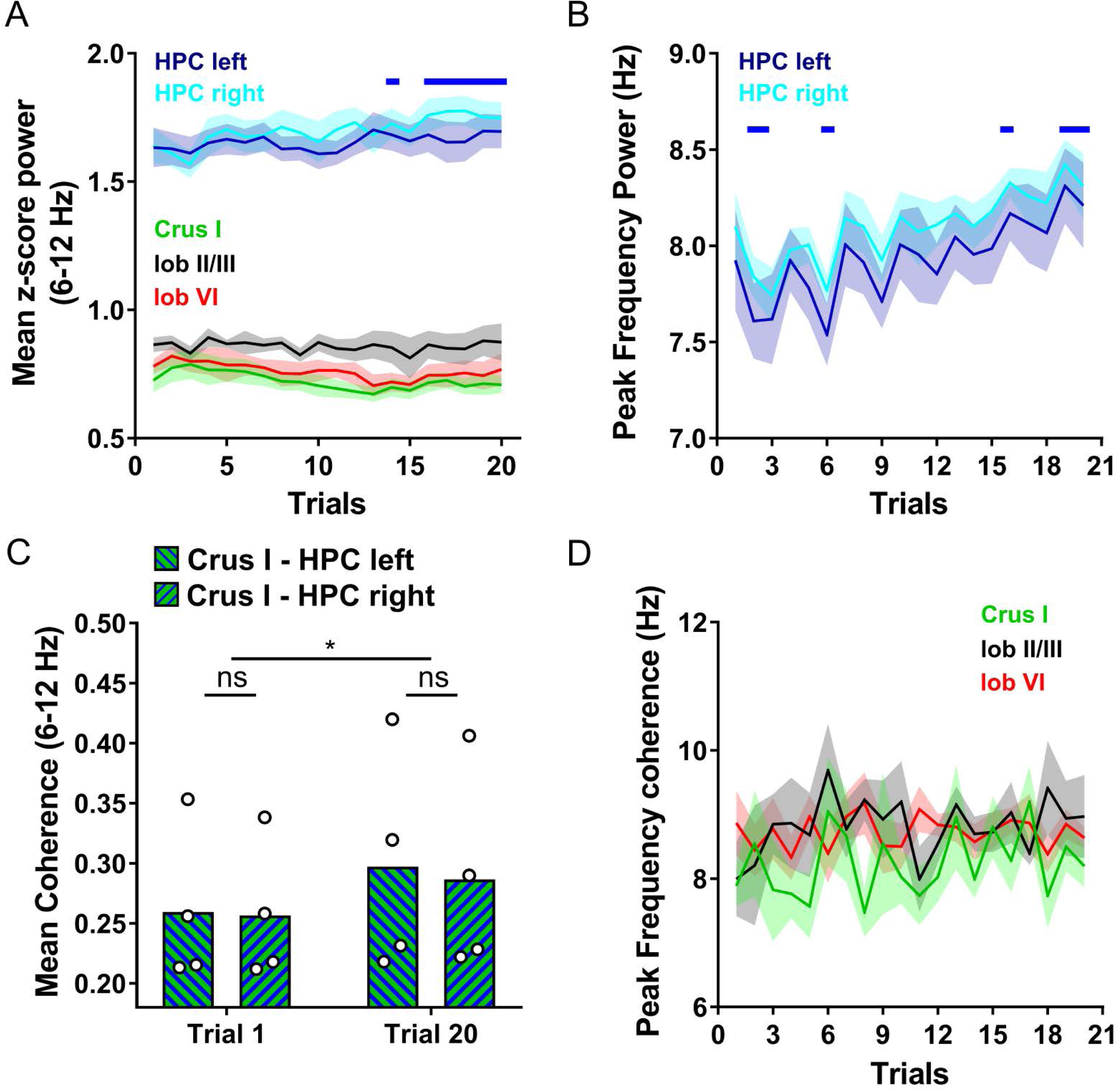
Cerebello-hippocampal coherence patterns are conserved across hemispheres during running on the linear track. **A**, Mean z-score 6-12 Hz power of the recorded LFPs from left (n = 6) and right (n = 5) hippocampus (HPC) and from the cerebellum (colour coded; Crus I, n = 5; lobule II/III, n = 6; lobule VI, n = 7) across trials. No laterality effect was found in the HPC (hemisphere x trial two-way ANOVA with FDR correction, hemisphere effect *p* = 0.5974, interaction effect *p* = 0.3132, trial effect *p* < 0.0001). Solid blue line indicate trials where HPC values were significantly higher than those in trial 1 (q < 0.05). **B**, Over trials, there was a significant increase in the peak frequency of the power spectra in both left and right HPC recordings; however, no differences were found between hemispheres (hemisphere x trial two-way ANOVA with FDR correction, trial effect p < 0.0001, hemisphere effect p = 0.4124, interaction effect p > 0.9999; solid blue line shows trials significantly different from trial 1, q < 0.05). **C**, No differences were observed in the mean 6-12 Hz coherence between left or right HPC and Crus I at early (trial 1) or late (trial 20) stages of training (Crus I-HPC left, n = 4; Crus I-HPC right, n = 4; hemisphere x trial two-way ANOVA, hemisphere effect p = 0.9026, interaction effect p = 0.7272, trial effect p = 0.0183). **D**, Peak frequency of cerebello-hippocampal coherence was not affected by the hippocampal hemisphere and did not change across trials for any combination (colour coded; Crus I-HPC left, n = 4, Crus I-HPC right, n = 4, hemisphere x trial two-way ANOVA, hemisphere effect p = 0.3601, interaction effect p = 0.9652, trial effect p = 0.2954; lobule II/III-HPC left, n = 4, lobule II/III-HPC right, n = 5, hemisphere x trial two-way ANOVA, hemisphere effect p = 0.2485, interaction effect p = 0.5048, trial effect p = 0.1767; lobule VI-HPC left, n = 6, lobule VI-HPC right, n = 6, hemisphere x trial two-way ANOVA, hemisphere effect p = 0.7025, interaction effect p = 0.9446, trial effect p = 0.2543). Shading indicates S.E.M.

**Figure S8.**
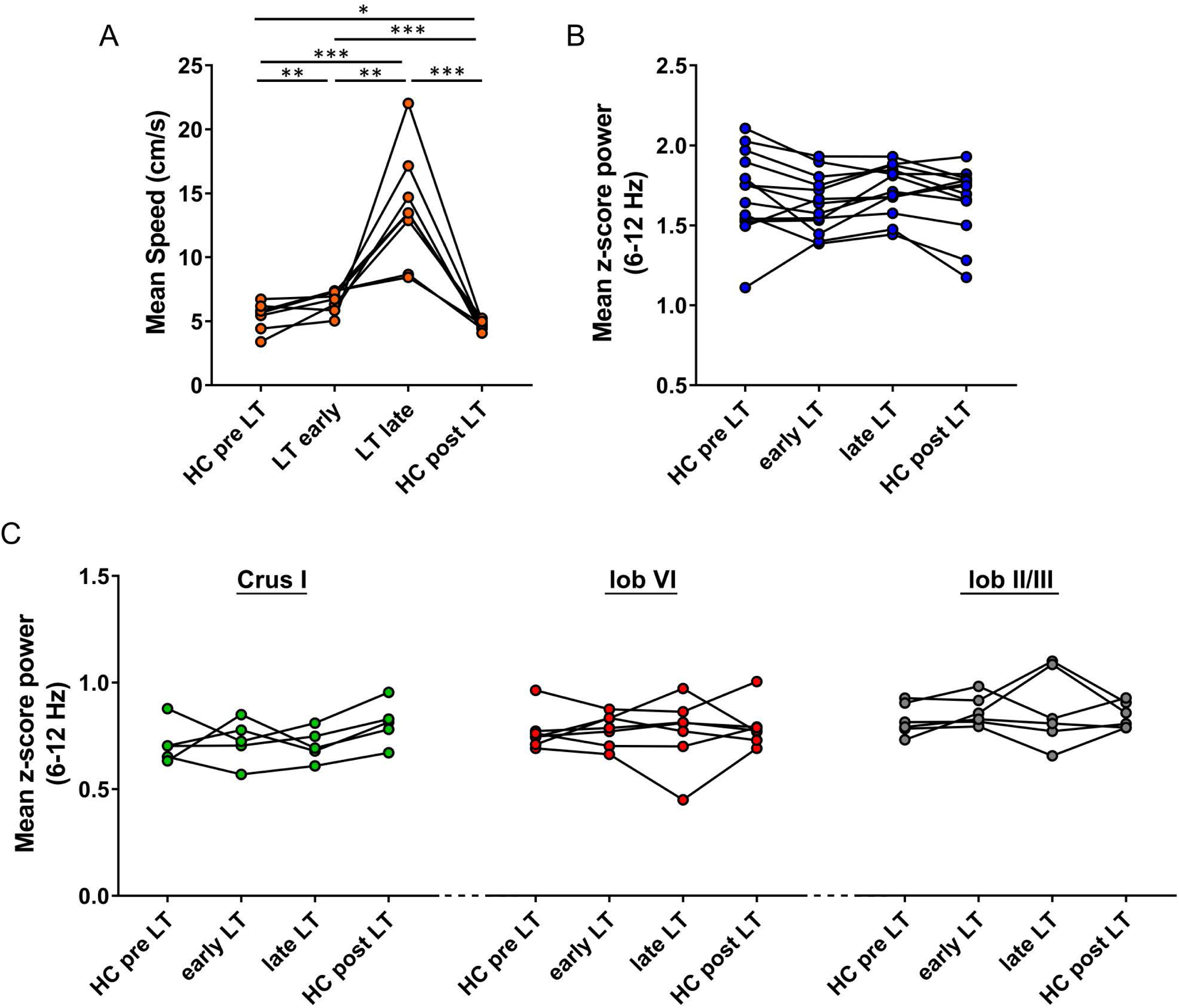
Speed and power spectrum dynamics across homecage and real world linear track conditions. **A,** Significant changes in speed were found across all conditions (HC pre LT, early and late LT and HC post LT; one-way repeated measures ANOVA with FDR correction, n = 8 mice). **B**, No significant differences were observed in hippocampal 6-12Hz z-score power (pooled values from both hemispheres) across conditions (repeated measures Friedman test, p = 0.1764, n = 13). **C**, No significant differences were observed across conditions in 6-12 Hz z-score power of Crus I (repeated measures Friedman test, p = 0.0755, n = 5), lobule VI (repeated measures Friedman test, p = 0.4188, n = 7) or lobule II/III (repeated measures Friedman test, p = 0.3751, n = 6). * q < 0.05, ** q < 0.01, *** q < 0.001. Abbreviations as for Fig. 5.

**Figure S9.**
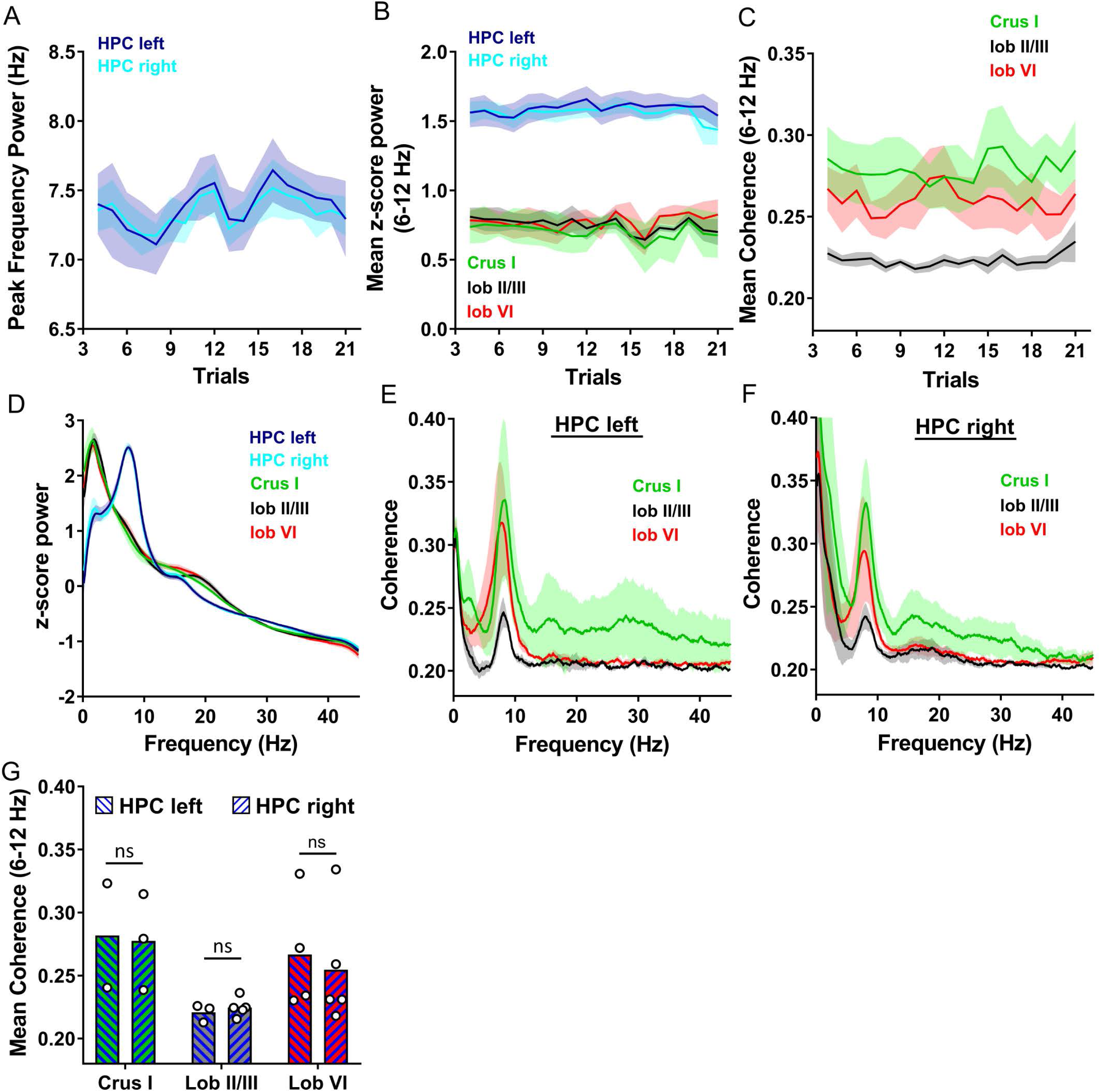
Cerebello-hippocampal coherence patterns are similar across hemispheres during goal-directed behavior in virtual reality. **A**, Peak frequency of the power spectra from left (n = 4) and right (n = 6) hippocampus (HPC) across trials. There was no difference between hemispheres but there was a significant effect of the trial (hemisphere x trial two-way ANOVA with FDR correction, trial effect p = 0.004, hemisphere effect p = 0.8795, interaction effect p = 0.9998). **B**, Mean z-score 6-12 Hz LFP power (colour coded; Crus I, n = 3; lobule II/III, n = 5; lobule VI, n = 5) across trials. No laterality effect or trial effect was found in the HPC (hemisphere x trial two-way ANOVA with FDR correction, hemisphere effect *p* = 0.7357, interaction effect *p* = 0.9865, trial effect *p* = 0.4180). Similarly, no differences were observed between cerebellar regions and no changes across trials (cerebellar region x trial two-way ANOVA with FDR correction, cerebellar region effect p = 0.8290, interaction effect p = 0.9993, trial effect p = 0.1781). **C**, Pooled, mean 6-12 Hz cerebello-hippocampal coherence (colour coded; Crus I-HPC = 5 values/3mice, lobule II/III = 8 values/5 mice, lobule VI-HPC = 9 values/5mice) across trials in the virtual reality condition. No differences across trials were observed at any cerebello-hippocampal combination (cerebellar region x trial two-way ANOVA with FDR correction, trial effect p = 0.0724, interaction effect p = 0.1265). **D**, Overall z-score power spectrum from cerebellar cortical regions (colour coded) and bilateral HPC averaged across all trials. **E**, Overall coherence between cerebellar cortical regions (Crus I, n = 2; lobule II/III, n = 3; lobule VI, n = 4; colour coded) and left HPC averaged across all VR trials. **F**, Overall coherence between cerebellar cortical regions (Crus I, n = 3; Lobule II/III, n = 5; Lobule VI, n = 5; colour coded) and right HPC. **G**, Mean 6-12 Hz coherence between cerebellar recordings and left or right HPC. No differences were observed between hemispheres (hemisphere x combination one-way ANOVA with FDR correction, hemisphere effect p = 0.6355). Shading indicates S.E.M.

## References

1. Petrosini L, Leggio MG, Molinari M. The cerebellum in the spatial problem solving: a co-star or a guest star? Prog Neurobiol. 1998;56: 191–210. Available: http://www.ncbi.nlm.nih.gov/pubmed/9760701

2. Rondi-Reig L, Burguière E. Is the cerebellum ready for navigation? Progress in brain research. 2005. pp. 199–212. doi:10.1016/S0079-6123(04)48017-0

3. Burguière E, Arleo A, Hojjati M reza, Elgersma Y, Zeeuw CI De, Berthoz A, et al. Spatial navigation impairment in mice lacking cerebellar LTD: a motor adaptation deficit? Nat Neurosci. 2005;8: 1292–1294. doi:10.1038/nn1532

4. Koziol LF, Budding D, Andreasen N, d’Arrigo S, Bulgheroni S, Imamizu H, et al. Consensus Paper: The Cerebellum’s Role in Movement and Cognition. The Cerebellum. 2014;13: 151–177. doi:10.1007/s12311-013-0511-x

5. Buckner RL. The Cerebellum and Cognitive Function: 25 Years of Insight from Anatomy and Neuroimaging. Neuron. 2013;80: 807–815. doi:10.1016/j.neuron.2013.10.044

6. Stoodley CJ, d’Mello AM, Ellegood J, Jakkamsetti V, Liu P, Nebel MB, et al. Altered cerebellar connectivity in autism and cerebellar-mediated rescue of autism-related behaviors in mice. Nat Neurosci. 2017;20: 1744–1751. doi:10.1038/s41593-017-0004-1

7. Kelly RM, Strick PL. Cerebellar loops with motor cortex and prefrontal cortex of a nonhuman primate. J Neurosci. 2003;23: 8432–44. Available: http://www.ncbi.nlm.nih.gov/pubmed/12968006

8. Ramnani N. The primate cortico-cerebellar system: anatomy and function. Nat Rev Neurosci. Nature Publishing Group; 2006;7: 511–522. doi:10.1038/nrn1953

9. Watson TC, Becker N, Apps R, Jones MW. Back to front: cerebellar connections and interactions with the prefrontal cortex. Front Syst Neurosci. 2014;8: 4. doi:10.3389/fnsys.2014.00004

10. Watson TC, Jones MW, Apps R. Electrophysiological mapping of novel prefrontal - cerebellar pathways. Front Integr Neurosci. 2009;3: 18. doi:10.3389/neuro.07.018.2009

11. Stoodley CJ, Schmahmann JD. Evidence for topographic organization in the cerebellum of motor control versus cognitive and affective processing. Cortex. 2010;46: 831–844. doi:10.1016/j.cortex.2009.11.008

12. Kim SG, Uğurbil K, Strick PL. Activation of a cerebellar output nucleus during cognitive processing. Science. 1994;265: 949–51. Available: http://www.ncbi.nlm.nih.gov/pubmed/8052851

13. Chen CH, Fremont R, Arteaga-Bracho EE, Khodakhah K. Short latency cerebellar modulation of the basal ganglia. Nat Neurosci. 2014;17: 1767–75. doi:10.1038/nn.3868

14. Rogers TD, Dickson PE, Heck DH, Goldowitz D, Mittleman G, Blaha CD. Connecting the dots of the cerebro-cerebellar role in cognitive function: Neuronal pathways for cerebellar modulation of dopamine release in the prefrontal cortex. Synapse. 2011;65: 1204–1212. doi:10.1002/syn.20960

15. Iglói K, Doeller CF, Paradis A-L, Benchenane K, Berthoz A, Burgess N, et al. Interaction Between Hippocampus and Cerebellum Crus I in Sequence-Based but not Place-Based Navigation. Cereb Cortex. 2015;25: 4146–4154. doi:10.1093/cercor/bhu132

16. Babayan BM, Watilliaux A, Viejo G, Paradis A-L, Girard B, Rondi-Reig L. A hippocampo-cerebellar centred network for the learning and execution of sequence-based navigation. Sci Rep. 2017;7: 17812. doi:10.1038/s41598-017-18004-7

17. Rochefort C, Lefort J, Rondi-Reig L. The cerebellum: a new key structure in the navigation system. Front Neural Circuits. 2013;7: 35. doi:10.3389/fncir.2013.00035

18. Yu W, Krook-Magnuson E. Cognitive Collaborations: Bidirectional Functional Connectivity Between the Cerebellum and the Hippocampus. Front Syst Neurosci. 2015;9: 177. doi:10.3389/fnsys.2015.00177

19. Iwata K, Snider R. Cerebello-hippocampal influences on the electroencephalogram. Electroencephalogr Clin Neurophysiol. 1959;11: 439–46. Available: http://www.ncbi.nlm.nih.gov/pubmed/13663818

20. Babb TL, Mitchell AG, Crandall PH. Fastigiobulbar and dentatothalamic influences on hippocampal cobalt epilepsy in the cat. Electroencephalogr Clin Neurophysiol. 1974;36: 141–154. doi:10.1016/0013-4694(74)90151-5

21. Snider RS, Maiti A. Septal afterdischarges and their modification by the cerebellum. Exp Neurol. 1975;49: 529–39. Available: http://www.ncbi.nlm.nih.gov/pubmed/811491

22. Krook-Magnuson E, Szabo GG, Armstrong C, Oijala M, Soltesz I. Cerebellar Directed Optogenetic Intervention Inhibits Spontaneous Hippocampal Seizures in a Mouse Model of Temporal Lobe Epilepsy. eNeuro. Society for Neuroscience; 2014;1. doi:10.1523/ENEURO.0005-14.2014

23. Rochefort C, Arabo A, Andre M, Poucet B, Save E, Rondi-Reig L. Cerebellum Shapes Hippocampal Spatial Code. Science (80-). 2011;334: 385–389. doi:10.1126/science.1207403

24. Rondi-Reig L, Paradis A-L, Lefort JM, Babayan BM, Tobin C. How the cerebellum may monitor sensory information for spatial representation. Front Syst Neurosci. 2014;8: 205. doi:10.3389/fnsys.2014.00205

25. Choe KY, Sanchez CF, Harris NG, Otis TS, Mathews PJ. Optogenetic fMRI and electrophysiological identification of region-specific connectivity between the cerebellar cortex and forebrain. Neuroimage. 2018;173: 370–383. doi:10.1016/j.neuroimage.2018.02.047

26. Arrigo A, Mormina E, Anastasi GP, Gaeta M, Calamuneri A, Quartarone A, et al. Constrained spherical deconvolution analysis of the limbic network in human, with emphasis on a direct cerebello-limbic pathway. Front Hum Neurosci. 2014;8: 987. doi:10.3389/fnhum.2014.00987

27. Newman PP, Reza H. Functional relationships between the hippocampus and the cerebellum: an electrophysiological study of the cat. J Physiol. 1979;287: 405–26. Available: http://www.ncbi.nlm.nih.gov/pubmed/430426

28. Whiteside J, Snider R. Relation of cerebellum to upper brain stem. J Neurophysiol. 1953;16: 397–413. Available: http://www.ncbi.nlm.nih.gov/pubmed/13070051

29. Snider RS, Maiti A. Cerebellar contributions to the papez circuit. J Neurosci Res. 1976;2: 133–146. doi:10.1002/jnr.490020204

30. Harper JW, Heath RG. Anatomic connections of the fastigial nucleus to the rostral forebrain in the cat. Exp Neurol. 1973;39: 285–92. Available: http://www.ncbi.nlm.nih.gov/pubmed/4573973

31. Heath RG, Dempesy CW, Fontana CJ, Myers WA. Cerebellar stimulation: effects on septal region, hippocampus, and amygdala of cats and rats. Biol Psychiatry. 1978;13: 501–29. Available: http://www.ncbi.nlm.nih.gov/pubmed/728506

32. Apps R, Hawkes R. Cerebellar cortical organization: a one-map hypothesis. Nat Rev Neurosci. 2009;10: 670–81. doi:10.1038/nrn2698

33. Fries P. A mechanism for cognitive dynamics: neuronal communication through neuronal coherence. Trends Cogn Sci. 2005;9: 474–480. doi:10.1016/j.tics.2005.08.011

34. Aoki S, Coulon P, Ruigrok TJH. Multizonal Cerebellar Influence Over Sensorimotor Areas of the Rat Cerebral Cortex. Cereb Cortex. 2017; doi:10.1093/cercor/bhx343

35. Suzuki L, Coulon P, Sabel-Goedknegt EH, Ruigrok TJH. Organization of cerebral projections to identified cerebellar zones in the posterior cerebellum of the rat. J Neurosci. 2012;32: 10854–69. doi:10.1523/JNEUROSCI.0857-12.2012

36. Mosko S, Lynch G, Cotman CW. The distribution of septal projections to the hippocampus of the rat. J Comp Neurol. 1973;152: 163–174. doi:10.1002/cne.901520204

37. Dolorfo CL, Amaral DG. Entorhinal cortex of the rat: topographic organization of the cells of origin of the perforant path projection to the dentate gyrus. J Comp Neurol. 1998;398: 25–48. Available: http://www.ncbi.nlm.nih.gov/pubmed/9703026

38. Witter MP. The perforant path: projections from the entorhinal cortex to the dentate gyrus. Progress in brain research. 2007. pp. 43–61. doi:10.1016/S0079-6123(07)63003-9

39. Coulon P, Bras H, Vinay L. Characterization of last-order premotor interneurons by transneuronal tracing with rabies virus in the neonatal mouse spinal cord. J Comp Neurol. 2011;519: 3470–3487. doi:10.1002/cne.22717

40. Brochu G, Maler L, Hawkes R. Zebrin II: a polypeptide antigen expressed selectively by Purkinje cells reveals compartments in rat and fish cerebellum. J Comp Neurol. 1990;291: 538–52. doi:10.1002/cne.902910405

41. Sugihara I, Shinoda Y. Molecular, topographic, and functional organization of the cerebellar cortex: a study with combined aldolase C and olivocerebellar labeling. J Neurosci. 2004;24: 8771–85. doi:10.1523/JNEUROSCI.1961-04.2004

42. Sugihara I, Quy PN. Identification of aldolase C compartments in the mouse cerebellar cortex by olivocerebellar labeling. J Comp Neurol. 2007;500: 1076–92. doi:10.1002/cne.21219

43. Sugihara I. Compartmentalization of the Deep Cerebellar Nuclei Based on Afferent Projections and Aldolase C Expression. The Cerebellum. 2011;10: 449–463. doi:10.1007/s12311-010-0226-1

44. Voogd J, Barmack NH. Oculomotor cerebellum. Progress in brain research. 2006. pp. 231–268. doi:10.1016/S0079-6123(05)51008-2

45. Ruigrok TJH. Ins and outs of cerebellar modules. Cerebellum. 2011;10: 464–74. doi:10.1007/s12311-010-0164-y

46. Cooke P, Snider R. Some cerebellar influences on electrically-induced cerebral seizures. Epilepsia. 1955;4: 19–28. Available: http://www.ncbi.nlm.nih.gov/pubmed/13305547

47. Jwair S, Coulon P, Ruigrok TJH. Disynaptic Subthalamic Input to the Posterior Cerebellum in Rat. Front Neuroanat. Frontiers; 2017;11: 13. doi:10.3389/fnana.2017.00013

48. Ugolini G. Advances in viral transneuronal tracing. J Neurosci Methods. 2010;194: 2–20. doi:10.1016/j.jneumeth.2009.12.001

49. Kelly RM, Strick PL. Rabies as a transneuronal tracer of circuits in the central nervous system. J Neurosci Methods. 2000;103: 63–71. doi:10.1016/S0165-0270(00)00296-X

50. Zheng Y, Goddard M, Darlington CL, Smith PF. Long-term deficits on a foraging task after bilateral vestibular deafferentation in rats. Hippocampus. 2009;19: 480–6. doi:10.1002/hipo.20533

51. Goddard M, Zheng Y, Darlington CL, Smith PF. Locomotor and exploratory behavior in the rat following bilateral vestibular deafferentation. Behav Neurosci. 2008;122: 448–59. doi:10.1037/0735-7044.122.2.448

52. Stackman RW, Clark AS, Taube JS. Hippocampal spatial representations require vestibular input. Hippocampus. 2002;12: 291–303. doi:10.1002/hipo.1112

53. Akaike T. The tectorecipient zone in the inferior olivary nucleus in the rat. J Comp Neurol. 1992;320: 398–414. doi:10.1002/cne.903200311

54. Teune TM, van der Burg J, van der Moer J, Voogd J, Ruigrok TJH. Topography of cerebellar nuclear projections to the brain stem in the rat. Progress in brain research. 2000. pp. 141–172. doi:10.1016/S0079-6123(00)24014-4

55. Herrero L, Yu M, Walker F, Armstrong DM, Apps R. Olivo-cortico-nuclear localizations within crus I of the cerebellum. J Comp Neurol. 2006;497: 287–308. doi:10.1002/cne.20976

56. Mihailoff GA, Burne RA, Azizi SA, Norell G, Woodward DJ. The pontocerebellar system in the rat: An HRP study. II. Hemispheral components. J Comp Neurol. 1981;197: 559–577. doi:10.1002/cne.901970403

57. Edge AL, Marple-Horvat DE, Apps R. Lateral cerebellum: functional localization within crus I and correspondence to cortical zones. Eur J Neurosci. 2003;18: 1468–85. Available: http://www.ncbi.nlm.nih.gov/pubmed/14511327

58. Glickstein M, Sultan F, Voogd J. Functional localization in the cerebellum. Cortex. 2011;47: 59–80. doi:10.1016/j.cortex.2009.09.001

59. Singer W. Neuronal synchrony: a versatile code for the definition of relations? Neuron. 1999;24: 49–65, 111–25. Available: http://www.ncbi.nlm.nih.gov/pubmed/10677026

60. De Zeeuw CI, Hoebeek FE, Schonewille M. Causes and consequences of oscillations in the cerebellar cortex. Neuron. 2008;58: 655–8. doi:10.1016/j.neuron.2008.05.019

61. Cheron G, Márquez-Ruiz J, Dan B. Oscillations, Timing, Plasticity, and Learning in the Cerebellum. The Cerebellum. 2016;15: 122–138. doi:10.1007/s12311-015-0665-9

62. Dugué GP, Brunel N, Hakim V, Schwartz E, Chat M, Lévesque M, et al. Electrical Coupling Mediates Tunable Low-Frequency Oscillations and Resonance in the Cerebellar Golgi Cell Network. Neuron. 2009;61: 126–139. doi:10.1016/j.neuron.2008.11.028

63. d’Angelo E, Nieus T, Maffei A, Armano S, Rossi P, Taglietti V, et al. Theta-frequency bursting and resonance in cerebellar granule cells: experimental evidence and modeling of a slow k+-dependent mechanism. J Neurosci. 2001;21: 759–70. Available: http://www.ncbi.nlm.nih.gov/pubmed/11157062

64. Hartmann MJ, Bower JM. Oscillatory activity in the cerebellar hemispheres of unrestrained rats. J Neurophysiol. 1998;80: 1598–604. Available: http://www.ncbi.nlm.nih.gov/pubmed/9744967

65. Wang Y, Chen H, Hu C, Ke X, Yang L, Xiong Y, et al. Baseline theta activities in medial prefrontal cortex and deep cerebellar nuclei are associated with the extinction of trace conditioned eyeblink responses in guinea pigs. Behav Brain Res. 2014;275: 72–83. doi:10.1016/j.bbr.2014.08.059

66. Rowland NC, Goldberg JA, Jaeger D. Cortico-cerebellar coherence and causal connectivity during slow-wave activity. Neuroscience. 2010;166: 698–711. doi:10.1016/j.neuroscience.2009.12.048

67. Chen H, Wang Y, Yang L, Sui J, Hu Z, Hu B. Theta synchronization between medial prefrontal cortex and cerebellum is associated with adaptive performance of associative learning behavior. Sci Rep. 2016;6: 20960. doi:10.1038/srep20960

68. Courtemanche R, Lamarre Y. Local Field Potential Oscillations in Primate Cerebellar Cortex: Synchronization With Cerebral Cortex During Active and Passive Expectancy. J Neurophysiol. 2004;93: 2039–2052. doi:10.1152/jn.00080.2004

69. Frederick A, Bourget-Murray J, Chapman CA, Amir S, Courtemanche R. Diurnal influences on electrophysiological oscillations and coupling in the dorsal striatum and cerebellar cortex of the anesthetized rat. Front Syst Neurosci. 2014;8: 145. doi:10.3389/fnsys.2014.00145

70. Soteropoulos DS, Baker SN. Cortico-Cerebellar Coherence During a Precision Grip Task in the Monkey. J Neurophysiol. 2005;95: 1194–1206. doi:10.1152/jn.00935.2005

71. O’Connor SM, Berg RW, Kleinfeld D. Coherent Electrical Activity Between Vibrissa Sensory Areas of Cerebellum and Neocortex Is Enhanced During Free Whisking. J Neurophysiol. 2002;87: 2137–2148. doi:10.1152/jn.00229.2001

72. Colgin LL. Mechanisms and Functions of Theta Rhythms. Annu Rev Neurosci. 2013;36: 295–312. doi:10.1146/annurev-neuro-062012-170330

73. Wikgren J, Nokia MS, Penttonen M. Hippocampo–cerebellar theta band phase synchrony in rabbits. Neuroscience. 2010;165: 1538–1545. doi:10.1016/j.neuroscience.2009.11.044

74. Onuki Y, Van Someren EJW, De Zeeuw CI, Van der Werf YD. Hippocampal–Cerebellar Interaction During Spatio-Temporal Prediction. Cereb Cortex. 2015;25: 313–321. doi:10.1093/cercor/bht221

75. Kajikawa Y, Schroeder CE. How Local Is the Local Field Potential? 2011; doi:10.1016/j.neuron.2011.09.029

76. Basso MA, May PJ. Circuits for Action and Cognition: A View from the Superior Colliculus. Annu Rev Vis Sci. 2017;3: 197–226. doi:10.1146/annurev-vision-102016-061234

77. Proville RD, Spolidoro M, Guyon N, Dugué GP, Selimi F, Isope P, et al. Cerebellum involvement in cortical sensorimotor circuits for the control of voluntary movements. Nat Neurosci. 2014;17: 1233–1239. doi:10.1038/nn.3773

78. Cerminara NL, Apps R, Marple-Horvat DE. An internal model of a moving visual target in the lateral cerebellum. J Physiol. 2009;587: 429–442. doi:10.1113/jphysiol.2008.163337

79. Cerminara NL, Apps R. Behavioural significance of cerebellar modules. Cerebellum. 2011;10: 484–94. doi:10.1007/s12311-010-0209-2

80. Iseni F, Lafay F, Raux H, Blondel D. Mapping of monoclonal antibody epitopes of the rabies virus P protein. J Gen Virol. 1997;78: 119–124. doi:10.1099/0022-1317-78-1-119

81. Franklin K, Paxinos G. The mouse brain in stereotaxic coordinates. 3rd ed. Elsevier; 2007.

82. de Lavilléon G, Lacroix MM, Rondi-Reig L, Benchenane K. Explicit memory creation during sleep demonstrates a causal role of place cells in navigation. Nat Neurosci. Nature Research; 2015;18: 493–495. doi:10.1038/nn.3970

83. Carlezon WA, Chartoff EH. Intracranial self-stimulation (ICSS) in rodents to study the neurobiology of motivation. Nat Protoc. 2007;2: 2987–2995. doi:10.1038/nprot.2007.441

84. Lasztóczi B, Klausberger T. Hippocampal Place Cells Couple to Three Different Gamma Oscillations during Place Field Traversal. Neuron. 2016;91: 34–40. doi:10.1016/j.neuron.2016.05.036

85. Bokil H, Andrews P, Kulkarni JE, Mehta S, Mitra PP. Chronux: A platform for analyzing neural signals. J Neurosci Methods. 2010;192: 146–151. doi:10.1016/j.jneumeth.2010.06.020

86. Islam MK, Rastegarnia A, Nguyen AT, Yang Z. Artifact characterization and removal for in vivo neural recording. J Neurosci Methods. 2014;226: 110–123. doi:10.1016/j.jneumeth.2014.01.027

